# Genome-wide association studies of seed metabolites identify loci controlling specialized metabolites in *Arabidopsis thaliana*

**DOI:** 10.1101/2022.09.23.509130

**Authors:** Thomas Naake, Federico Scossa, Leonardo Perez de Souza, Monica Borghi, Yariv Brotman, Tetsuya Mori, Ryo Nakabayashi, Takayuki Tohge, Alisdair R. Fernie

## Abstract

Plants synthesize specialized metabolites to facilitate environmental and ecological interactions. During evolution, plants diversified in their potential to synthesize these metabolites. Quantitative differences in metabolite levels of natural *Arabidopsis thaliana* accessions can be employed to unravel the genetic basis for metabolic traits using genome-wide association studies (GWAS). Here, we performed metabolic GWAS (mGWAS) on seeds of a panel of 315 *A. thaliana* natural accessions, including the reference genotypes C24 and Col-0, for polar and semi-polar seed metabolites using untargeted ultra-performance liquid chromatography-mass spectrometry. As a complementary approach, we performed quantitative trait locus (QTL) mapping of near-isogenic introgression lines between C24 and Col-0 for specific seed specialized metabolites. Besides common QTL between seeds and leaves, GWAS revealed seed-specific QTL for specialized metabolites indicating differences in the genetic architecture of seeds and leaves. In seeds, aliphatic methylsulfinylalkyl and methylthioalkyl glucosinolates associated with the *GS-ALK* and *GS-OHP* locus on chromosome 4 containing *alkenyl hydroxyalkyl producing 2* (*AOP2*) and *3* (*AOP3*) and/or with the *GS-ELONG* locus on chromosome 5 containing *methylthioalkyl malate synthase* (*MAM1*) and *MAM3*. We detected two unknown sulfur-containing compounds that were also mapped to these loci. In GWAS, some of the annotated flavonoids (kaempferol 3-*O*-rhamnoside-7-*O*-rhamnoside, quercetin 3-*O*-rhamnoside-7-*O*-rhamnoside) were mapped to *transparent testa 7* (*AT5G07990*), encoding a cytochrome P450 75B1 monooxygenase. Three additional mass signals corresponding to quercetin-containing flavonols were mapped to *UGT78D2* (*AT5G17050*). The association of the loci and associating metabolic features were functionally verified in knockdown mutant lines. By performing GWAS and QTL mapping, we were able to leverage variation of natural populations and parental lines to study seed specialized metabolism. The GWAS data set generated here is a high-quality resource that can be interrogated in further studies.

## Introduction

Two main phenotypic novelties have been critical during the transition from an aquatic to a terrestrial environment. The first of these innovations was the emergence of phenylpropanoid and lignin biosynthesis, allowing early terrestrial plants to acquire a relatively rigid body structure and colonize the land (Weng and Chapple, 2010). The second innovation consisted in the development of structures specialized for reproduction and dispersal, like pollen and seeds. These were essential for long-distance transport and successful colonization of the new environment by the offspring of primordial land plants (Linkies et al., 2010; Willis et al., 2014). Seeds, as a reproductive structure, also needed to be protected from adverse environmental conditions, including fungal attacks, insect feeding, or UV radiation. The chemical composition of seeds was thus selected not only to provide the essential nutrients during germination, but also to accumulate a number of specialized metabolites conferring protective properties against biotic and abiotic stresses (Debeaujon et al., 2000).

*Arabidopsis thaliana* is an ideal model to study the link between phenotypic and genomic variation, given the wealth of genomic resources available (Alonso-Blanco et al., 2016; Togninalli et al., 2018). The considerable genetic variation of Arabidopsis was employed to study local adaptation in collections of natural accessions (Seren et al., 2017). GWAS is a technique to leverage natural variation and was used in previous studies to detect adaptive traits (Atwell et al., 2010; Togninalli et al., 2018). GWAS assesses the effect of each genomic marker at the population level, represented by information on high-density SNPs, on a quantitatively-assessed phenotype with the likelihood of the association (Seren et al., 2017). QTL mapping, in comparison to GWAS, identifies genomic regions that co-segregate with a given trait in lines resulting from biparental or lately also multiparental crosses. GWAS and QTL mapping were employed to study primary metabolism (Chan et al., 2010a; Wu et al., 2016; Slaten et al., 2020), specialized metabolism (Kliebenstein et al., 2001a; Hansen et al., 2008; Chan et al., 2010b; Chan et al., 2011; Routaboul et al., 2012; Li et al., 2014; Bac-Molenaar et al., 2015; Ishihara et al., 2016; Tohge et al., 2016; Wu et al., 2018), heavy metal (Chao et al., 2012) and salt tolerance (Baxter et al., 2010), shade avoidance (Filiault and Maloof, 2012), and flowering time (Li et al., 2010).

In *A. thaliana*, two major classes of specialized metabolites have been considered to confer protective properties to abiotic stress, namely flavonoids and glucosinolates. Flavonoids are arguably the best-characterized class of specialized metabolites that are universally distributed in the plant kingdom (Winkel-Shirley, 2001; Winkel-Shirley, 2002; Falcone Ferreyra et al., 2012; Tohge et al., 2017). By analyzing flavonoid-less mutants (*ban*, *ap2*, and *transparent testa*), Debeaujon et al. (2000) could show that a lack of flavonoids resulted in lower dormancy and structural aberrations in seeds (missing layers, modified epidermal layers). Tepfer et al. (2012) and Tepfer and Leach (2017) showed that flavonoid-less seed mutants exhibited lower survival rate when exposed to solar UV and cosmic radiation for 1.5 years. Generally, besides being involved in developmental and photoprotective processes, flavonoids convey antioxidative properties (Seyoum et al., 2006; Mierziak et al., 2014) and play a role in biotic stress (Treutter, 2005; Mierziak et al., 2014); as yet, however, no such information is available if the same is true in seeds. In *A. thaliana* seeds, a wide array of flavonoids can be found, mainly belonging to the subclasses of flavonols (mono- and diglycosylated quercetin, kaempferol, and isorhamnetin derivatives) and of flavan-3-ols (epicatechin monomers and procyanidin polymers, Routaboul et al., 2006).

The other major class of specialized metabolites conferring tolerance to abiotic stress, glucosinolates, are mainly restricted to the Brassicales order, including the Brassicaceae, Capparaceae, and Caricaceae families, but were also found in at least 500 non-cruciferous angiosperm species (Fahey et al., 2001). The glucosinolate biosynthetic pathway and its regulation is well studied (Supplementary Figure S1, Kliebenstein et al., 2001a; Kliebenstein et al., 2001b; Grubb and Abel, 2006; Halkier and Gershenzon, 2006; Hirai et al., 2007; Seo and Kim, 2017). Glucosinolates are mainly attributed to be involved in biotic stress response (Grubb and Abel, 2006; Halkier and Gershenzon, 2006; Samuni-Blank et al., 2012). The role of glucosinolates in stress response was mainly defined through functional analysis of overexpression lines or mutants deficient in their regulation or biosynthesis (Beekwilder et al., 2008; Zhang et al., 2015). In *A. thaliana* seeds, 34 different glucosinolate species were detected that revealed different accumulation patterns in 39 different Arabidopsis ecotypes (Kliebenstein et al., 2001a). Two major glucosinolate subclasses, methylthioalkyl and methylsulfinylalkyl glucosinolates, showed striking differences between accessions: while the accessions Bs-1, Aa-0, Ma-0 and Yo-0 showed high methylthioalkyl:methylsulfinylalkyl glucosinolates ratio in seeds (> 5), 13 accessions showed a ratio > 3 (e.g., Sei-0, Tsu-1, and Mrk-0, Kliebenstein et al., 2001a). Furthermore, Kliebenstein et al. (2001a) found that glucosinolate accumulation differs between leaves and seeds: (*i*) the accessions Kas and Sorbo accumulate low levels of 2-hydroxy-3-butenyl glucosinolate in leaves, but high levels of this glucosinolate in seeds; (*ii*) the methylthioalkyl:methylsulfinylalkyl glucosinolates ratio in seeds is for all accessions > 1, while for leaves this was only found in three accessions (Bla-10, Can-0, Su-0).

Previous studies in our group revealed differences in seed glucosinolate levels of *A. thaliana* Col-0, C24 (unpublished data) in introgression lines (Törjék et al., 2008). Taken together with previous findings that showed differences in the accumulation of seed specialized metabolites in Arabidopsis ecotypes, we conducted an untargeted metabolic profiling analysis by UPLC coupled to high-resolution mass spectrometry (MS) on *A. thaliana* seed polar and semi-polar metabolites (covering several classes of specialized metabolites) to reveal quantitative differences of metabolites between accessions. To find putative novel gene candidates that control the accumulation of specialized metabolites, we conducted GWAS and, in a complementary approach, QTL mapping on the Arabidopsis IL population obtained from the cross between C24 and Col-0. We show here that (*i*) previously characterized metabolites (flavonoids and glucosinolates) associate with known loci, (*ii*) two unknown sulfur-containing metabolites map to glucosinolates-associated loci, and (*iii*) that the respective Arabidopsis SALK knockdown lines of the gene *AT5G17050*, previously selected from GWAS, showed quantitative changes in the levels of the associated quercetin-containing flavonol compounds.

## Results and Discussion

### Genome-wide association studies of untargeted seed metabolite analysis shows a large set of mass feature pairs associated with the same loci

Genetic natural variation is an indispensable resource to find genes that are involved in the biosynthesis and regulation of plant specialized metabolites (Matsuda et al., 2015; Chen et al., 2016). Here, we determined the relative levels of polar and semipolar seed metabolites from about 300 *A. thaliana* ecotypes using UPLC-MS from two growing seasons (replicate 1 and replicate 2) and from one previously published set of leaf metabolites (Wu et al., 2018), and mapped the features to their associated genomic loci using the same GWAS approach we applied previously (Zhu et al., 2022). This approach encompasses mixed linear models to account for the amount of phenotypic covariance caused by the genetic relatedness, which should reduce confounding effects due to the population structure and kinship (Yu et al., 2006; Kang et al., 2008; Zhang et al., 2010; Vilhjálmsson and Nordborg, 2013). Due to computational constraints, we did not identify epistatic interactions, even though these will contribute to the observed phenotypes (Marchini et al., 2005; Cordell, 2009; Kam-Thong et al., 2011; Chen et al., 2014; Dong et al., 2015; Kerwin et al., 2015). Epistasis is the interaction of genetic variation at multiple loci that results in non-additive effects in the analyzed phenotypes (Soltis and Kliebenstein, 2015). In Arabidopsis, epistatic interactions typically involve the interaction of three or more loci (Wentzell et al., 2007; Rowe et al., 2008; Joseph et al., 2013a; Joseph et al., 2013b; Kerwin et al., 2015).

To compare metabolites across the different sets, we matched the alignments of mass features of seed replicate 1 and seed replicate 2/leaf based on their *m/z* deviation, retention time deviation, and the covariance between the two seed replicates resulting in 21007 features for the negative and 36194 features for the positive ionization mode. To further refine the accuracy, we imposed stricter matching rules, adjusting for retention time shift between replicate 1 and replicate 2 and a correlation of > 0.1, resulting in a total number of 9008 features for the negative and 12133 features for positive ionization mode (core set). 2882 (negative ionization mode) and 3798 (positive ionization mode) matched mass features, i.e. those conserved between the aligned replication datasets, were mapped to the same locus/loci in GWAS (Supplementary Figure S2). Those features that were mapped to the same locus/loci generally had higher heritability values (H^2^, Supplementary Figure S1 A, negative ionization mode: all: 0.549, mapped: 0.666; positive ionization mode: all: 0.542, mapped: 0.665) and higher correlation values (Supplementary Figure S1 B, negative ionization mode: all: 0.408, mapped: 0.534; positive ionization mode: all: 0.401, mapped: 0.529) than random pairs from the complete core set.

In a next step, we created a table with the GWAS results of the two biological replicates of seeds and of the leaf samples. We reported for each joint mass feature the assigned QTL and LOD scores. From the core set, the two replicates from seed GWAS showed high positive Spearman correlation values for both data sets acquired in negative (⍴ = 0.485) and positive (⍴ = 0.497) ionization mode. The Spearman correlation values were lower (⍴ between 0.181 and 0.269) when comparing the seed replicates with the result from leaf GWAS (Figure 1 A and B). Furthermore, when looking at the intersection of shared loci (Figure 1 C), we found that shared loci between the seed replicates showed a higher number (4884 for negative and 5688 for positive ionization mode) compared to that between seed and leaf GWAS (2242/1332 for negative and 2935/1942 for positive ionization mode). This may indicate that the reproducibility between the seed replicates of the core set is higher as when compared to the results from leaf GWAS. Alternatively, this may reflect a degree of variation in the genetic architecture. The comparison of the seeds and leaf datasets allowed the identification of tissue-specific QTL (Wu et al., 2018) and highlighted the different genetic architecture of these two tissues in controlling the accumulaton of some specific mass features. However, there were 1026 and 1247 loci controlling the mass features in the core sets that are shared between the two seed replicates and the leaf data set, indicating conserved loci controlling the levels of mass features across different tissues. The mass feature pairs that showed overlap between the two seed replicates, and for some of these also to the leaf data set, represents a highly valuable resource that we make fully available in the Supplemental Material. In the subsequent paragraphs we only focus on those associations related to glucosinolates and flavanols, as well as on some unknown mass features supposedly representing novel glucosinolates and flavonoids.

**Figure 1:**
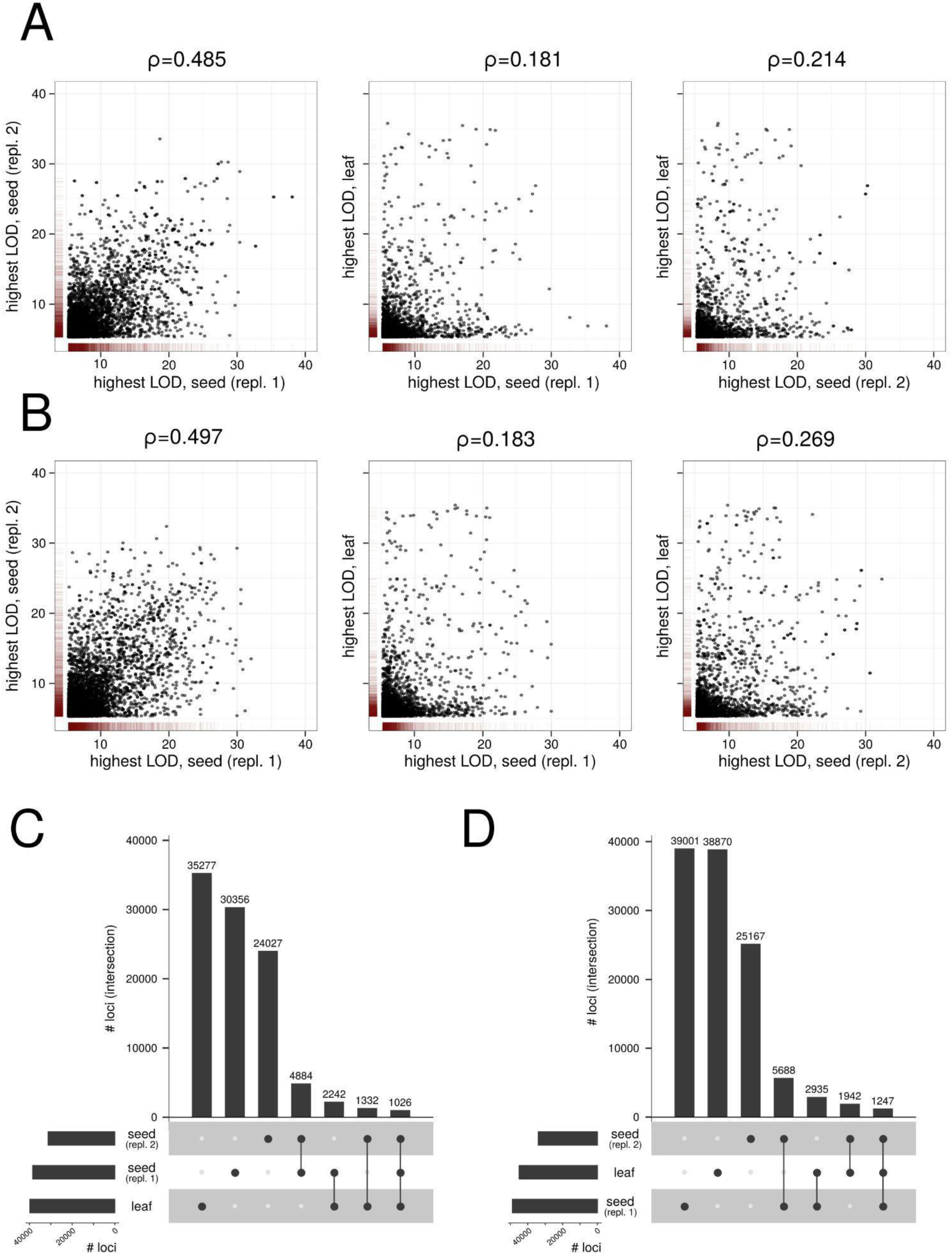
Mapping of seed replicates 1 and 2 and leaf in genome-wide association studies. A and B: Scatterplot of highest LOD values of shared QTL per matched mass feature pair for negative (A) or positive ionization mode (B). The colored lanes display the density of data points. The Spearman correlation ⍴ values are indicated for the different sets. C and D: Intersection sets of QTL for mass feature pairs with LOD > 5.3 for negative (C) and positive ionization mode (D). LOD: logarithm of odds; QTL: quantitative trait loci.

Alongside the shared loci, the majority of loci were not shared between the different mass feature sets (Figure 1 C and D). When analyzing the distribution of the proportion of mass features mapped to loci, the sets that did not show intersection (seed replicate 1, seed replicate 2, leaf) had a higher proportion of two or more of mass features mapped to the same loci compared to sets that show intersection (Supplementary Figure S3). This could be attributed to measurement errors, associations of non-causative markers with a given trait, driven by linkage with causative markers (Korte and Farlow, 2013), reflect environmental variance, and/or genotype-environment interaction effects. We assume that the significant associations from the intersection sets (Figure 1 C and D), being conserved between the different replicates, represent genuine QTL characterized by lower sources of error.

As a complementation, we performed QTL mapping using biparental NILs from Col-0 and C24 (Törjék et al., 2008). This population was useful in detecting saiginols (phenylacylated flavonols) in floral tissues (Tohge et al., 2016). In the GWAS population and NIL population of Arabidopsis seeds, saiginols were not detected.

### Variation in glucosinolate levels in seeds is controlled by the *GS-ELONG*, *GS-ALK*, and *GS-OHP* loci

Based on previous studies, we annotated several metabolites in our data set, including amino acids, glucosinolates, and metabolites from the flavonoid and phenylpropanoid biosynthetic pathways (Supplementary Table S5 and S6). The annotation of glucosinolates included all known methylsulfinylalkyl and methylthioalkyl glucosinolates. Most of these metabolites showed high broad-sense heritability (H^2^ > 0.75) and were mostly mapped with high LODs for both replicates to a locus on chromosome 5 and, for some of these glucosinolate metabolites, to a locus on chromosome 4 (Figure 2 A and Supplementary Figure S4 A for 3-hydroxypropyl glucosinolate, Supplementary Table S7 and S8). Within the locus on chromosome 5, the genes *methylthioalkylmalate synthase 1* (*MAM1*, *AT5G23010*) and *3* (*MAM3*, *AT5G23020*) are located, previously named the *GS-ELONG* locus. *MAM1* catalyzes the condensation reaction of two cycles of chain elongation in methionine-derived glucosinolate biosynthesis and a *mam1* mutant showed a decrease in C_4_ and an increase in C_3_ glucosinolates (Kroymann et al., 2001). MAM3 accepts all ⍵-methylthio-2-oxoalkanoic acids required to synthesize C_5_ to C_8_ aliphatic glucosinolates in *A. thaliana* (Textor et al., 2007). Within the locus on chromosome 4, the genes *AOP1* (*AT4G03070*), *AOP2* (*AT4G03060*), and *AOP3* (*AT4G03050*) are located, which are known as the *GS-ALK* and *GS-OHP* locus. The *AOP* genes encode 2-OG dependent dioxygenases that are involved in glucosinolate biosynthesis. AOP2 and AOP3 convert methylsulfinylalkyl glucosinolates into alkenyl glucosinolates and hydroxyalkyl glucosinolates, respectively (Kliebenstein et al., 2001b; Kliebenstein et al., 2001a). The parental lines of the NIL population, C24 and Col-0, of the NIL population showed strong differences in relative glucosinolate levels: 4-methylsulfinylbutyl glucosinolate showed 8.40-times and 4-methylthiobutyl glucosinolate 5.05-times higher levels in Col-0 compared to C24; 3-butenyl glucosinolate showed 23.5-times and 8-methylsulfinyloctyl glucosinolate 265-times higher levels in C24 compared to Col-0. Subsequently, aliphatic glucosinolates showed strong relative metabolite differences in QTL mapping for genomic regions containing Col-0 *AOP* (Supplementary Figure S5, Col-0 *AOP* in MASC05042-MASC09225 referring to the lines M36, M20, and M21).

**Figure 2:**
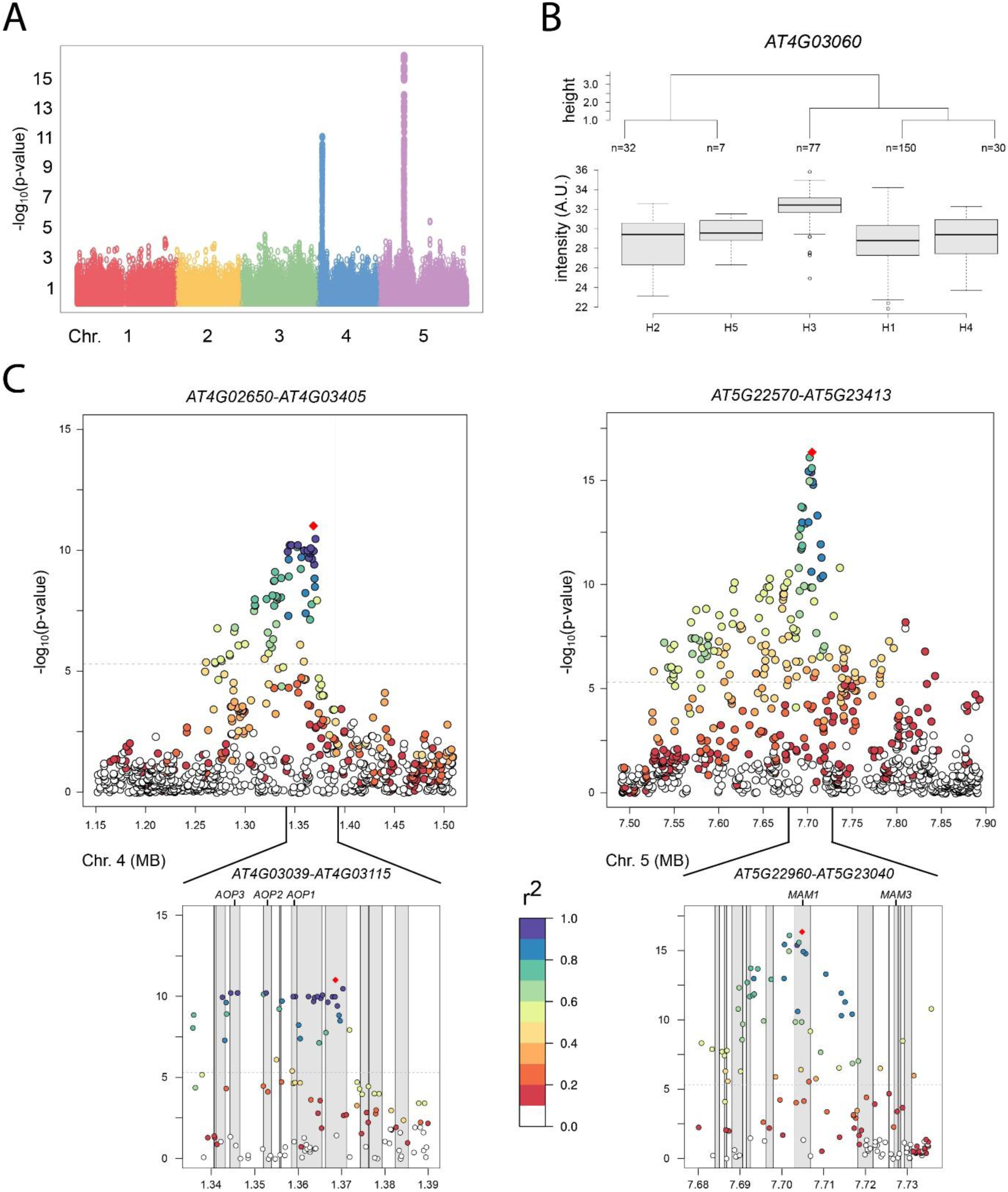
Genome-wide association mapping for 3-hydroxypropyl glucosinolate (negative ionization mode). A: The Manhattan plot of 3-hydroxypropyl glucosinolate shows two peaks in each replicate on chromosomes 4 (highest LOD: 11.01) and 5 (16.35). These loci contain the genes *AOP1*, *AOP2*, and *AOP3* (chromosome 4), *MAM1* and *MAM3* (chromosome 5) that are involved in glucosinolate biosynthesis. Only replicate 1 is shown here. B: Haplotype analysis of metabolite levels of 3-hydroxypropyl glucosinolate. The nucleotide sequence differences were statistically associated with the levels of 3-hydroxypropyl glucosinolate (ANOVA q-value: 1.78e-20 for replicate 1). Only data for replicate 1 is shown in A and B. The data for replicate 2 is depicted in Supplementary Figure S4. C: LD analysis of the associated genomic regions on chromosome 4 and 5 for 3-hydroxypropyl glucosinolate. The locus on chromosome 4 shows LD for the genomic region containing the genes *AOP1*, *AOP2*, and *AOP3*, while the locus on chromosome 5 marks a sharper decrease in standardized LD (r^2^) indicating that the *MAM1* gene is mainly responsible for the natural diversity of 3-hydroxypropyl glucosinolate levels. *AOP*: *alkenyl hydroxyalkyl producing*; A.U.: arbitrary units; LD: linkage disequilibrium; LOD: logarithm of odds; *MAM*: *methylthioalkylmalate synthase*.

Haplotypes of the genes *MAM1*, *MAM3*, and *AOP1*, *AOP2*, and *AOP3* showed significant differences in metabolite levels according to ANOVA (Figure 2 B and Supplementary Figure S4 B for 3-hydroxypropyl glucosinolate, GWAS population) indicating that the allelic variation at these target loci is responsible for the observed metabolite differences. Indeed, some of the SNPs for these genes involved in glucosinolate biosynthesis resulted in amino acid differences (Supplementary Table S9). The LD analysis for 3-hydroxypropyl glucosinolate revealed that the alleles on chromosome 4 are in high LD (standardized LD r^2^ close to 1) spanning the genomic region containing *AOP1*, *AOP2*, and *AOP3* (Figure 2 C, left panel). In *A. thaliana*, LD usually decays 50% within 5 kb (Gan et al., 2011; Korte and Farlow, 2013). Here, the loci containing *AOP1*, *AOP2*, and *AOP3* showed wider LD. The situation on chromosome 5 marks a sharp decrease for 3-hydroxypropyl glucosinolate and peaks in the gene region of *MAM1* (*AT5G23010*). Interestingly, *MAM3* was in high LD (r^2^ > 0.6) with the SNP showing the highest LOD in *MAM1*, but did not show as high r^2^ values as neighboring genes within the locus on chromosome 4. This indicates that *MAM1* is the main gene controlling 3-hydroxypropyl glucosinolate levels. Previously, these loci were also detected from GWAS of glucosinolate levels in leaves (Chan et al., 2011). The Arabidopsis *gtr1 gtr2* double mutant, which lacks (or contains low amounts, depending here on the type of the mutations it carries) the nitrate/peptide transporters responsible for glucosinolate transport to seeds, did not accumulate glucosinolates in seeds and exhibited a tenfold over-accumulation in the source tissues leaves and silique walls (Nour-Eldin et al., 2012). Thus, it seems that the variation in glucosinolate levels is ‘inherited’ from these source tissues.

### Unknown sulfur-containing metabolites map to *GS-ELONG*, *GS-ALK*, and/or *GS-OHP* loci in genome-wide association mapping

Next to the annotated glucosinolates, other mass features in the core set also showed association with the *GS-ELONG*, *GS-ALK*, and/or *GS-OHP* loci in GWAS. In particular, two mass features with *m/z* of 596.1104 (unknown 596) and 626.1032 (unknown 626) were mapped to chromosome 4 or 5. The unknown 626 was mapped to the GS-ELONG locus (both seed replicates and leaf had a LOD ≥ 5.3 for *GS-ELONG* locus, Supplementary Figure S6), while the unknown 596 was mapped for both seed replicates to the GS-ALK and GS-OHP loci (LOD ≥ 5.3). Correlated mass features that showed *m/z* differences defined by the transformations (Supplementary Table S4) also showed associations to these loci (Supplementary Figure S7).

The LD analysis revealed that for the unknown 626 the SNP with the highest LOD was located near or within the *MAM1* gene. The standardized LD, r^2^, decreased sharply when moving away from the *MAM1* gene (Supplementary Figure S6). To reveal the chemical composition of the two unknowns, we fed isotope-labeled ^13^C and ^34^S to the siliques and analyzed the metabolites by LC-QTOF-MS. The unknown 596 (*m/z* 596.1104 in negative ionization mode) and 626 (*m/z* 626.1032 in negative ionization mode) contain most probably 20 C atoms and 22 C atoms, respectively, based on isotope feeding experiments with ^13^C. The MS analysis indicated for the feeding experiments with ^34^S that the two unknown compounds contain two S atoms (Supplementary Figure S8). Interestingly, the QTL mapping between C24 and Col-0 introgression lines of the unknowns 596 and 626 identified an additional locus close, but not overlapping, to *GS-ALK* and *GS-OHP* (*AT4G15733*-*AT4G24620*, M50_2_ in Supplementary Figure S5). From the GWAS analysis here, the candidate region for the unknown 596 is *AT4G00005*-*AT4G03770*, overlapping with *AOP1*, *AOP2*, and *AOP3*. The introgression lines M20, M21, and M36 (Supplementary Figure S5 B), corresponding to the region *AT4G02465*-*AT4G08280*, showed higher levels of the unknowns 596 and 626 compared to the C24 background (Supplementary Figure S5 A). Thus, it is unclear if the *GS-ELONG*, *GS-ALK*, and *GS-OHP* loci directly control the levels of the unknowns 596 and 626 or if, by an indirect effect, *AOP* and *MAM* change the flux of sulfur-containing metabolites.

### Untargeted genome-wide association mapping of non-annotated mass features identifies a gene controlling flavonoid levels

The mass features with *m/z* 463.0885/465.1032 (7.08 min), 593.1516/595.1670 (6.73 min), 609.1466/611.1617 (6.20 min, negative/positive ionization mode, *m/z* values and retention time from the UPLC-MS analysis of SALK lines), and co-eluting mass features were mapped to the genes *AT5G17040* and *AT5G17050* (*UGT78D2*, Figure 3 A and Supplementary Figure S9) in GWAS. Given the characteristic *m/z* of 303.0504 and other spectrometric peaks (positive ionization mode), these metabolites were putatively annotated as quercetin hexoside (*m/z* 463.0885/465.1032), quercetin deoxyhexoside deoxyhexoside (*m/z* 593.1516/595.1670), and quercetin hexoside deoxyhexoside (*m/z* 609.1466/611.1617). The chromatographic peaks were distinct from other flavonols with the same *m/z*, e.g., quercetin 3-*O*-rhamnoside-7-*O*-rhamnoside (retention time of replicate 1: 6.71 min, unknown 593.1516/595.1670: 6.43 min).

**Figure 3:**
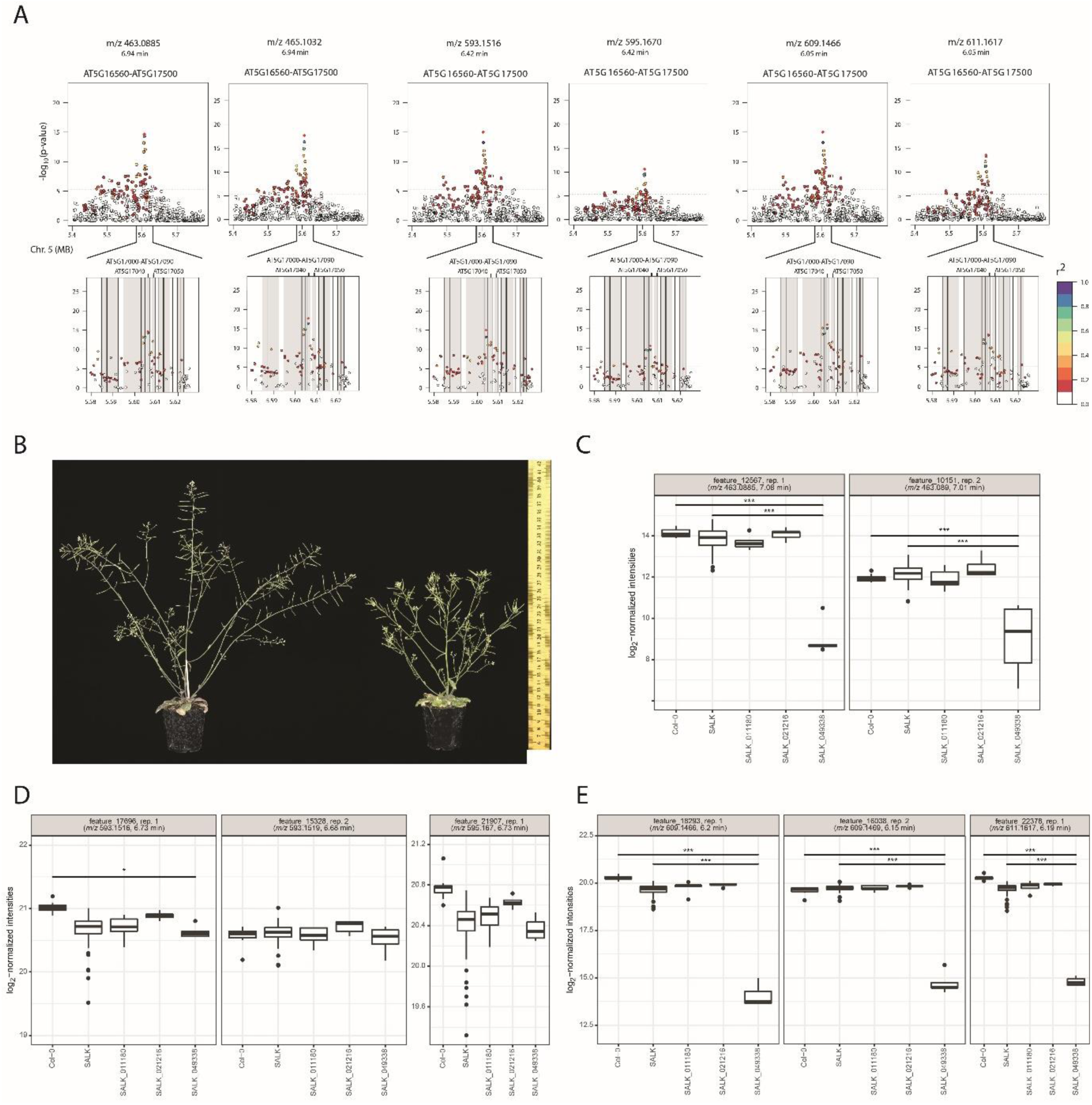
Flavonoid biosynthetic pathway: mutant phenotype and functional analysis. A: Linkage disequilibrium analysis for quercetin-containing flavonols (negative and positive ionization mode). The highest LOD is achieved for a SNP within the region of *AT5G17030* or *AT5G17040*. Standardized LD r^2^ is relatively low for the SNPs that are located within the gene *AT5G17050*. Only data for replicate 1 is shown in A. The data for replicate 2 is depicted in Supplementary Figure S9. B: The mutant of *AT5G17050* (SALK_049338) exhibited a dwarf phenotype with a loss of apical dominance (stunted inflorescence) as reported previously by Yin et al. (2014). C-E: Metabolite analysis of mapped mass features 463.0885/465.1032, 593.1516/595.1670, and 609.1466/611.1617 (negative/positive ionization mode) showed lower levels in the seeds of mutant lines (n = 5 individual plants) compared to wild-type Col-0 (n = 9). A.U.: arbitrary units. LD: linkage disequilibrium; LOD: logarithm of odds; MB: megabase; SNP: single nucleotide polymorphism. *: p-value < 0.05, **: p-value < 0.01, ***: p-value < 0.001

To validate the associations, we selected genes of interest based on (*i*) the LOD score from GWAS; (*ii*) the expression of the gene from the data reported by Schmid et al. (2005) (Affymetrix ATH1 array); (*iii*) haplotype and LD analysis, and (*iv*) potential involvement of the gene in the biosynthetic pathway based on homology analysis and literature support for quercetin-containing flavonols and other non-annotated mass features. For the flavonol-related metabolites, we selected two genes of interest; for unknown mass features in our core set, we selected seven genes of interest, and obtained T-DNA insertion SALK lines for functional validation. Except for three SALK lines, which showed to be heterozygous for the insertion, homozygosity was confirmed by PCR genotyping in the T2 generation (Supplementary Table S10) and SALK lines were individually cultivated in two replicates. Seeds of the SALK lines were analyzed by UPLC-MS (Supplementary Figure S10). The resulting data set was analyzed in terms of presence and differential abundance of the associated mass features with respect to Col-0 and other SALK line seeds in negative and positive ionization mode. Only the line SALK_049338 (*AT5G17050*, encoding UDP-GLUCOSYL TRANSFERASE 78D2) showed differential abundance for several mass features compared to the control lines (Supplementary Table S11 and S12).

When growing the line SALK_049338 (*AT5G17050*), we observed shorter stature for all plants as compared to the wild type (Figure 3 A and Supplementary Figure S11), a phenotype also previously reported when mutating this gene (Yin et al., 2014). The quercetin-containing flavonols 463.0885/465.1032 and 609.1466/611.1617 in this line exhibited lower seed metabolite levels compared to the other SALK lines (excluding the lines for *AT5G17040* and *AT5G17050*) and wild-type Col-0, while levels of 593.1516/595.1670 were not affected by *AT5G17050* (Figure 3 B).

Flavonoids are involved in the regulation of auxin transport (Buer and Muday, 2004; Peer and Murphy, 2007). Lee et al. (2005) and Tohge et al. (2005) described that UGT78D2 is a flavonoid 3-*O*-glucosyltransferase and that *ugt78d2* mutants show an altered flavonoid pattern. A *ugt78d1* (*AT1G30530*) *ugt78d2* double mutant exhibited a strong and specific repression of flavonol biosynthesis and was strongly impaired in the initial 3-*O*-glycosylation, while UGT78D3 (AT5G17030) only contributed to a minor extent to overall 3-*O*-glycosylation (Jones et al., 2003; Tohge et al., 2005; Yonekura-Sakakibara et al., 2008; Yin et al., 2012).

UGT73C6 (AT2G36790) is the 7-*O*-glucosyltransferase in flowers; however, 7-*O*-rhamnosylation by UGT89C1 (AT1G06000) is more common as the form of 7-*O*-conjugation (Yonekura-Sakakibara et al., 2008). Yin et al. (2014), studying *UGT78D2*, suggested that kaempferol 3-*O*-rhamnoside-7-*O*-rhamnoside is responsible for the altered growth phenotype by narrowing down the potential active moieties using a series of mutants. In the same study, a *ugt78d1 ugt78d2* double mutant showed strongly reduced levels of kaempferol 3-*O*-glucoside-7-*O*-rhamnoside and kaempferol 3-*O*-[rhamnosyl (1→2 glucoside)]-7-*O*-rhamnoside, while kaempferol 3-*O*-rhamnoside-7-*O*-rhamnoside was not detected at all. Furthermore, the levels of the aglycones kaempferol and quercetin were reduced to 21 % and 18 % of the wild-type levels, respectively.

Interestingly, the unknown quercetin deoxyhexoside deoxyhexoside (*m/z* 593.1516/595.1670), presumably containing rhamnoside, did not show lower levels in the *ugt78d2* mutant lines, despite the fact that the unknown flavonol showed association to *UGT78D2* in GWAS. This could be explained by the fact that *UGT78D2* is a glucosyltransferase, not a rhamnosyltransferase (Yin et al., 2014) and could indicate that *UGT78D2* indirectly controls the flux of rhamnosylated (deoxyhexosylated) flavonols in seeds. In our GWAS data set, kaempferol 3-*O*-rhamnoside-7-*O*-rhamnoside (H^2^ = 0.837 in positive ionization mode) showed association with a gene in the region *AT5G01680*-*AT5G13170*, but not with the locus containing *AT5G17050* (Supplementary Table S8). The SNP with the highest LOD (> 7.8 in positive ionization mode) located close to *transparent testa 7* (*tt7*, *AT5G07990*, data not shown). TT7 is a cytochrome P450 75B1 monooxygenase, an enzyme previously reported to have 3’-flavonoid hydroxylase activity (Schoenbohm et al., 2000) that regulates the kaempferol/quercetin ratio (Peer et al., 2001). Similarly, quercetin 3-*O*-rhamnoside-7-*O*-rhamnoside (H^2^ = 0.921 in positive ionization mode) was mapped with a LOD > 6.2 close to *TT7* (data not shown, positive ionization mode). On the other hand, kaempferol 3-*O*-glucoside-7-*O*-rhamnoside (H^2^ = 0.173 in positive ionization mode) had its highest LOD within the gene *UGT78D3* (replicate 2, no mapping with LOD > 5.3 for replicate 1). For the other annotated flavonol glycosides (in positive ionization mode) in the core set, no genome-wide association was obtained. For QTL mapping, no associations with flavonoids were detected. This is to be expected, since annotated flavonoid levels in the biparental lines showed little differences: kaempferol 3-*O*-glucoside-7-*O*-rhamnoside showed 1.24-times, quercetin 3-*O*-glucoside-7-*O*-rhamnoside 1.10-times, kaempferol 3-*O*-rhamnoside 1.26-times, and quercetin 3-*O*-rhamnoside 1.16-times higher levels in C24 compared to Col-0 (Supplementary Figure S5). For GWAS, missing associations could be due to low absolute variation of these metabolites or because these flavonoids are regulated by multiple loci that are not reported as significant in our approach. Higher differences in accumulation patterns can be triggered through application of different kinds of stress (e.g., UV radiation) before analyzing metabolite levels. This finding was generally in line with a previous smaller scale study that detected quantitative rather than qualitative differences in flavonoids between *A. thaliana* accessions and concluded that most flavonoids are controlled by a few additive loci with relatively broad effects (Routaboul et al., 2012).

Here, we focused on the analysis of the association involving candidate structural genes. In this paper we focused on candidate structural genes of the glucosinolate/flavonoid pathways, although we report in the Supplementary Material the full list of significant associations that may represent a resource to investigate the additional control these metabolites may have at the level of pathway regulation. Furthermore, the results from glucosinolates and some of the flavonoids indicated pleiotropic effects and collocating QTL for joint mass features. This analysis can be extended to a wider scale and to non-biosynthetic enzymes. Moreover, the core set exhibited differences in QTL between the mass features from the two seed replicates and the leaf replicate. Future studies will investigate the variation in the genetic architecture of traits controlling the levels of specialized metabolites across different tissues.

## Conclusion

Here, we performed GWAS on metabolic mass features of two biologically independent replicates of seed from two growing seasons and one replicate of leaves obtained by untargeted UPLC-MS. As a complementary approach, we performed QTL mapping of NIL introgression lines between C24 and Col-0 for specific seed metabolites. By including GWAS of leaf metabolites, we detected 4884 and 5688 loci for mass feature pairs (negative and positive ionization mode) that were exclusively detected for seed GWAS, indicating differences in tissue-specific associations between seeds and leaves. On the other hand, 1026 and 1247 QTL for mass feature pairs (negative and positive ionization mode) were conserved across seed and leaf tissues in GWAS.

In seeds, aliphatic methylsulfinylalkyl and methylthioalkyl glucosinolates as well as two unknown sulfur-containing compounds, tentatively identified as novel glucosinolates, showed associations in GWAS and QTL mapping with the known *GS-ELONG*, *GS-ALK*, and/or *GS-OHP* loci. In addition, QTL mapping detected an adjacent region on chromosome 4 for the two unknown sulfur-containing compounds. In GWAS, some of the annotated flavonoids in seeds showed associations to regions containing *TT7* or *UGT78D2*, including three previously unknown quercetin-containing flavonols. QTL mapping did not reveal any association for flavonoids. This difference is potentially caused by the low allelic variance in flavonoid-biosynthetic genes resulting in small differences in flavonoid levels in the parental lines.

A SALK knockdown line of the gene *UGT78D2* (*AT5G17050*) showed decreased levels of the quercetin-containing flavonols, while SALK lines of the neighboring gene *AT5G17040* did not show changes in flavonol levels. We would like to draw the following conclusions regarding the genetic architecture of seed specialized metabolism: (*i*) seed specialized metabolism differs substantially from leaf metabolism as shown by the identification of QTL that differ between these tissues, but the two tissues also exhibit common genetics to some degree; (*ii*) *AOP* and *MAM* genes are key regulators for glucosinolate seed metabolite levels in seeds. Aliphatic glucosinolates are presumably not synthesized *in situ* in seeds, but are transported from source tissues to seeds. The variation of aliphatic glucosinolates is ‘inherited’ from these source tissues; (*iii*) the alleles of *UGT78D2* (*AT5G17050*) affect the levels of quercetin-containing flavonols in seeds. The natural GWAS population was shaped by processes of genetic adaptation and meiotic events during evolution. This results in greater phenotypic variance compared to the NIL population between Col-0 and C24 as exemplified by differences in flavonoid levels. However, the overlap suggests, as previously stated (Brog et al., 2019), that genome-wide association and QTL mapping are complementary techniques to study seed specialized metabolism.

## Materials and Methods

### Plant material

The HapMap collection of natural *A. thaliana* accessions (315 accessions) with existing SNP data (Li et al., 2010; Horton et al., 2012) was used to perform GWAS on polar and semi-polar metabolites. Seed material for GWAS analysis was provided by Yariv Brotman (MPI-MP, Potsdam, Germany) and grown by the Green team of the Max Planck Institute of Molecular Plant Physiology in two growing seasons in the years 2017 (replicate 1) and 2018 (replicate 2) according to Wu et al. (2018). Seeds were sown directly to soil in 6 cm pots for each accession and stratified in a growth chamber (Percival Scientific, Perry, USA; 250 μE m^-2^ s^-1^ day/night 16 h/8 h, temperature of 20°C/6°C, relative humidity, RH, 60%/75%). After two weeks (end of March), the seedlings were pricked and transferred to separate pots with six replicates per accession. Plants were randomly placed in a polytunnel greenhouse (with an integrated frost protection system) and randomly dislocated every 1-2 weeks to avoid positional shading. Plants were bagged one month before harvest for seed collection (glassine bags, 40 g m^-2^). Two weeks before harvest, watering was stopped. Plants were harvested from the end of May until the middle of June depending on the genotype. Harvested bagged inflorescences were stored for four weeks at 15°C and 15% RH. Seeds were collected by sieving siliques (sieve size 355, Edinger Direkt, Leinburg, Germany) into glass vials before storing them at 15°C, 15% RH. Leaf samples for GWAS analysis were obtained by Wu et al. (2018) using the control condition samples.

The introgression line population of Arabidopsis (near-isogenic lines, NILs), obtained from the cross between Col-0 and C24 (Törjék et al., 2008), was cultivated as described in Tohge et al. (2016). Seeds were collected from three individual plants of 45 M lines (C24 background) and 69 N lines (Col-0 background) as described above.

The SALK lines (SALK_008908, SALK_011180, SALK_020876, SALK_021216, SALK_024438, SALK_027837, SALK_037430, SALK_049338, SALK_072964, SALK_081021, SALK_201809C, SALK_203337C, SALK_203919C, SALK_204674C, SALK_206494C) were obtained from the NASC database. C24, Col-0, and SALK mutant lines were cultivated under greenhouse conditions (21/19°C, day/night 16/8h, RH 50%/50%, additional illumination by Philips Son-T Agro lamps from 6 a.m.-10 a.m. and 6 p.m.-10 p.m.; Philips, Eindhoven, The Netherlands). The plants of the different lines were randomly placed to avoid block effects during growth. Plants were watered daily with 1/1000 Hyponex solution (Hyponex, Osaka, Japan). The trays with plants were randomly distributed two times per week to prevent positional light effects. Seeds were collected as described above.

### Genotyping of Col-0 and SALK lines

About 4 weeks after germination, one leaf per replicate (in total five replicates) was collected from Col-0, C24, and SALK lines, frozen in liquid N_2_, and stored at −80°C. DNA was extracted according to (Kasajima et al., 2004). Col-0 wild-type plants and the SALK lines were genotyped by PCR using the following mix: 15.7 μL water, 2 μL 10X DreamTaq buffer, 0.4 μL 10mM dNTP, 0.4 μL LBb1.3 or line-specific forward primer, 0.4 μL line-specific reverse primer, 0.1 μL DreamTaq polymerase (Thermo Fisher Scientific, Waltham, USA), 1 μL template DNA. The primers are described in Supplementary Table S1. The following program was used (Biometra T Professional Thermocycler, Analytik Jena, Jena, Germany): 5 min initial denaturation, 95°C; 35 cycles of 30 s denaturation, 95°C, 30 s annealing 58°C, 1 min extension, 72°C; 10 min final extension, 72°C; hold, 4°C. 20 μL PCR product were separated on a 1% agarose gel for 25 min at 120 V.

### Extraction of polar and semi-polar metabolites in seeds and leaves

Metabolites from seeds were extracted according to (Tohge and Fernie, 2010). 200 μL of pre-cooled (−20°C) 80% MeOH (Sigma-Aldrich, Munich, Germany; containing 1 µg isovitexin and 0.04 mg ribitol as internal standard) was added to 30 *A. thaliana* seeds (cooled in liquid N_2_), of which the weight was previously determined. After shaking the tubes, previously cooled in liquid N_2_ (3 min, 25 *Hz* by Retsch mill MM 301, Haan, Germany), the tubes were centrifuged for 10 min at room temperature (17,900 *g*), and the supernatant was transferred to a new tube. The tubes were centrifuged for 10 min at room temperature (17,900 *g*). 135 μL of the supernatant were transferred to a new tube, dried by speed-vac for 2-3 h, filled with argon, and stored at −80°C. On the day of analyses, the samples were resuspended in 100 μL 80% MeOH and transferred to sample vials.

Metabolites from leaves were extracted from 50 mg leaf material (cooled in liquid N_2_) using 500 μL of the same extraction buffer as above. The same extraction protocol was followed as above transferring 200 μL of the supernatant to a new tube before drying by speed-vac for 2-3 h. On the day of analyses, the samples were resuspended in 200 μL 80% MeOH and transferred to sample vials.

### Determination of relative polar and semi-polar metabolite levels by UPLC-MS for genome-wide association studies

For leaf and seed metabolites, extracts from Col-0, prepared as described above, were taken as a quality control. Metabolites were separated by Waters Acquity UPLC I using a Waters Acquity UPLC BEH C18 1.7 μm VanGuard^TM^ 2.1 x 5 mm as a pre-column and a Waters Acquity UPLC HSS T3 1.8 μm 2.1 x 100 mm as a column (Waters, Dublin, Ireland; injection volume 5 μL, sample temperature 10°C, column temperature 40°C, flow rate 0.4 mL min^-1^). The gradient was as follows: from 0 min to 1 min 99% buffer A (Water UL/C MS grade (Bio-Lab ltd., Jerusalem, Israel) + 0.1% formic acid) and 1% buffer B (100% acetonitrile UL/C MS grade (Bio-Lab ltd., Jerusalem, Israel) + 0.1% formic acid), 11 min 60% A and 40% B, 13 min 30% A and 70% B, 15 min 1% A and 99% B isocratic flow to 16 min, 17 min 99% A and 1% B isocratic flow to 20 min. Metabolites were ionized by ESI in negative and positive ionization mode(capillary voltage ±3.5 kV, sheath gas flow 60, auxiliary gas flow 20, capillary temperature 275°C, drying gas temperature 300°C, skimmer voltage 25 V, tube lens voltage 130 V). MS spectra were acquired from 1-20 min by Thermo Scientific Q Exactive in Full MS mode (resolution 70000, max. injection time 100 ms, automatic gain control value 3E6; Thermo Fisher Scientific, Waltham, USA) in the scan range 100-1500 *m/z*. Peaks per replicate and ionization mode were aligned by `Genedata` (version 10.5.3) using the settings according to Supplementary Table S2. Mass features that eluted before 0.5 min and after 16 min were removed from the peak alignment. For each ionization mode separately, the replicates were combined by matching based on a *m/z* deviance of ±0.01 and a retention time deviance of ±0.3 min to obtain the joint mass features present in both replicates. Intensity values were divided by the respective analyzed seed weight. Intensity values were log_2_ transformed and batch effects were removed by the function `removeBatchEffect` from the `limma` package (v3.38.3, Ritchie et al., 2015). In the case of multiple matches from replicate 1 to replicate 2, only the matched feature pairs with highest covariance are retained. Outliers were removed by checking their intensity values by boxplots and by projecting them via principal component analysis (PCA) by the function `prcomp`} from the `stats` package (v.4.1.2) in `R`.

### Determination of relative polar and semi-polar metabolite levels by HPLC-MS and QTL mapping

Metabolite levels were determined according to Tohge et al. (2016) using an HPLC system Surveyor (high pressure LC; Thermo Finnigan, Waltham, USA) coupled to a Finnigan LTQ-XP system (Thermo Finnigan, Waltham USA). Chromatographic data were processed via Xcalibur (v2.1, Thermo Fisher Scientific, Waltham, USA). QTL mapping was done according to Tohge et al. (2016).

### ^13^C and ^34^S isotope feeding and measurement by LC-quadrupole time-of-flight (QTOF) MS

*A. thaliana* seeds were labeled with ^13^C (via ^13^CO_2_) and ^34^S (via Na_2_^34^SO_4_) according to Nakabayashi et al. (2013) and Nakabayashi et al. (2016) using Col-0 plants prepared by SI Science Co., Ltd. (Saitama, Japan). The dried samples were extracted with 150 μl for ^13^C samples and 50 μl for ^34^S of 80% MeOH containing 2.5 μM 10-camphour sulfonic acid per mg dry weight using a mixer mill with zirconia beads for 7 min at 18 Hz and 4 C. After centrifugation for 10 min, the supernatant was filtered using an HLB μElution plate (Waters). The extracts (1 μl) were analyzed using LC-QTOF-MS (LC, Waters Acquity UPLC system; MS, Waters Xevo G2 Q-Tof). Analytical conditions were as follows LC: column, Acquity bridged ethyl hybrid (BEH) C18 (1.7 μm, 2.1 mm 100 mm, Waters); solvent system, solvent A (water including 0.1% [v/v] formic acid) and solvent B (acetonitrile including 0.1% [v/v] formic acid); gradient program, 99.5%A/0.5%B at 0 min, 99.5%A/0.5%B at 0.1 min, 20%A/80%B at 10 min, 0.5%A/99.5%B at 10.1 min, 0.5%A/99.5%B at 12.0 min, 99.5%A/0.5%B at 12.1 min and 99.5%A/0.5%B at 15.0 min; flow rate, 0.3 ml/min at 0 min, 0.3 ml/min at 10 min, 0.4 ml/min at 10.1 min, 0.4 ml/min at 14.4 min and 0.3 ml/min at 14.5 min; column temperature, 40 C; MS detection: polarity, negative; capillary voltage, −2.75 kV; cone voltage, 25.0 V; source temperature, 120 C; desolvation temperature, 450 C; cone gas flow, 50 l/h; desolvation gas flow, 800 l/h; collision energy, 6 V; mass range, m/z 50‒1500; scan duration, 0.1 sec; interscan delay, 0.014 sec; data acquisition, centroid mode; Lockspray (Leucine enkephalin); scan duration, 1.0 sec; interscan delay, 0.1 sec.

### Determination of relative polar and semi-polar metabolite levels by UPLC-MS for Col-0, C24, and SALK mutant lines

Metabolites were separated by Waters Acquity UPLC using a Waters HSS T3 C18 (Waters, Dublin, Ireland, 100 mm I. x 2.1 mm i.d. x 1.8 μm particle size) as column and pre-column (column temperature 40°C, flow rate 0.4 mL min^-1^). The gradient was as follows: 1 min 99% buffer A (Water UPLC MS grade + 0.1% formic acid; Biosolve, Dieuze, France) and 1% buffer B (100% acetonitrile + 0.1% formic acid; Biosolve, Dieuze, France), 11 min 60% A and 40 B, 13 min 30% A and 70% B, 15 min 1% A and 99% B isocratic flow to 16 min, 17 min 99% A and 1% B isocratic flow to 20 min. Metabolites were ionised by ESI in negative and positive ionisation mode (capillary voltage ±3 kV, sheath gas flow 60, auxiliary gas flow 35, capillary temperature 150°C, drying gas temperature 350°C, skimmer voltage 25 V, tube lens voltage 130 V). MS spectra were acquired from 1-19 min by ThermoScientific Q Exactive in MS mode (resolution 25000, max. injection time 100 ms, automatic gain control value 1E6; Thermo Fisher Scientific, Waltham, USA;) in the scan range 100-1500 *m/z*. Peaks were aligned by `xcms` (v3.16.1, Smith et al., 2006) and annotated by `CAMERA` (v.1.50.0, Kuhl et al., 2012) in the `R` programming language (v4.1.2, see Supplementary Table S3). Intensity values were divided by the respective seed weight. Intensity values were log_2_ transformed and batch effects were removed by the function `removeBatchEffect` from the `limma` package (v3.38.3). Outliers were removed by checking their quality via the MatrixQCvis package (v1.5.4, Naake and Huber, 2022). Metabolite and mass features were checked by the Thermo Xcalibur Qual Browser (v4.0.27.21, Thermo Fisher Scientifc, Waltham, USA).

### Genome-wide association mapping, calculation of heritability, haplotype and linkage disequilibrium analysis, and statistical testing for differences in SALK lines

A similar approach to Fusari et al. (2017) and Wu et al. (2018) was taken to map metabolite information to genetic loci. The `R` packages `EMMAX` (Efficient Mixed-Model Association eXpedited, Kang et al., 2010) and `GAPIT` (Genomic Association and Prediction Integrated Tool, version 23-May-18, Lipka et al., 2012) were used to perform the mapping. We employed a mixed linear model containing fixed and random effects and characterized the population structure using the first three principal components (Q matrix, Price et al., 2006) to incorporate this information together with the VanRaden kinship matrix (Eu-Ahsunthornwattana et al., 2014) as fixed and random effects, respectively (`method = “MLM”`). The aligned mass feature table with normalized intensity values was used as an input. The `GAPIT` function was used to map the phenotypic observations (normalized metabolite intensities) to loci in the *A. thaliana* genome using 199455 SNP markers with minor allele frequency > 1% obtained using Affymetrix GeneChip Array 6.0 (TAIR version 9, Li et al., 2010; Horton et al., 2012) using `PCA.total = 3`, `model = “MLM”`, `SNP.fraction = 1.0` (all other parameters were set to default). The logarithm of odds (LOD) threshold was set to 5.3 (-log_10_(1/*N*) with *N* the number of SNPs). The resulting SNPs with LOD ≥ 5.3 were assigned to the same group if the genomic distance between them was less than 10 kb and the genes within the respective groups were considered as candidate genes.

Broad-sense heritability (H^2^) was defined by the proportion of the total variance explained by the genetic variance according to Fusari et al. (2017) using the `lmer` function and obtaining the variances by the function `VarCorr` from `lme4` (v1.1-23, Bates et al., 2015). For calculating the heritability, only the features were used that showed a retention time deviance of ≤ 0.075 min (retention time_repl. 1_ - retention time_repl. 2_), an absolute *m/z* deviance of ≤ 0.075, and a Pearson correlation of > 0.1.

For haplotype analysis, the distance between haplotypes was calculated from the SNPs by the `dist.gene` function from the `ape` (v5.3, Paradis and Schliep, 2019) package (`method = “pairwise”`, `pairwise.deletion = FALSE`, `variance = FALSE`). Distances were clustered by the `hclust` function (`method = “ward.D”`) and the tree was cut by `cutree` (`h = 0.00001`) from the `stats` package (v3.6.2). To test for statistical relation between haplotypes and metabolite levels, ANOVA (`anova` from the `stats` package, v3.6.2) was performed with FDR correction (false discovery rate, `p.adjust` with `method = “BH”`), adjusting for the number of all metabolites used for mapping in negative and positive ionization mode. For linkage disequilibrium (LD) analysis, the p-values were taken from the GWAS results file for the respective mass feature (LOD = −log_10_(p-value)). Standardized LD, r^2^, values were calculated via the function `r2fast` from the `GenABEL` package (v1.8-0, Aulchenko et al., 2007). Expression analysis for genes of interest was conducted within the eFP browser (Winter et al., 2007) using the data set of Schmid et al. (2005).

To test for differences in SALK lines, the log2-normalized raw intensities were tested against Col-0 or the respective complement of SALK lines using limma (v3.50.3). To this end, linear models were fitted for each metabolic feature using `lmFit` and moderated t-statistics were computed by empirical Bayes moderation of the standard errors towards a global value using `eBayes` (`trend = TRUE`). p-values were adjusted using FDR via the Benjamini-Hochberg method. Since there was no corresponding second replicate available in positive ionization mode, the corresponding features in negative ionization mode were determined using correlation analysis and retention time window thresholding. If multiple features in negative ionization mode matched to the feature in positive ionization mode, the feature with highest correlation to the feature of positive mode (replicate 1) was selected.

The scripts can be found at https://www.github.com/tnaake/GWAS_arabidopsis_seed.

### MetNet network construction

*m/z* and retention time values of seed replicate 1 were used for structural network inference via `structural` and `rtCorrection` from the `MetNet` package (v1.15.3, `R` v4.1.2, Naake and Fernie, 2019) using the transformations and retention time shifts described in Supplementary Table S4. Edges corresponding to adduct additions were removed if the retention time between two mass features was > 0.1 min. The combined peaklists with log-normalized intensity values of replicate 1 and 2 were used as input for statistical network construction (function `statistical`) using Pearson and Spearman correlation. The weighted statistical adjacency matrices were thresholded (function `threshold`) only retaining correlation values > 0.7 for Pearson and Spearman correlation coefficients and FDR-adjusted p-values < 0.05 using the Benjamini-Hochberg method. The network was visualized in `Cytoscape` (v3.7.2, Shannon et al., 2003). The script can be found at https://www.github.com/tnaake/GWAS_arabidopsis_seed.

## Author Information

### Author contributions

T.N., Y.B., T.T., and A.R.F. designed experiments. T.N. and F.S. harvested plants and extracted metabolites. T.N., L.P.S., and Y.B. measured and processed metabolite data sets. T.N. performed GWAS mapping and downstream analysis. T.T. performed QTL mapping and downstream analysis. T.N. and T.T. annotated metabolites in the data set. R. N., T.M., and T.T. performed ^13^C-and ^34^S-feeding experiments and obtained the related data. T.N. and M.B. performed genotyping for SALK lines. T.N. analyzed metabolites of SALK lines. T.N. and A.R.F. wrote the manuscript with input from all other authors.

### Notes

The authors declare no competing financial interest.

## Supporting information

Supplemental_Material_loci_negative_mode

Supplemental_Material_loci_positive_mode

## Acknowledgements

We would like to thank Elena Doubijanski, Ben-Gurion University of the Negev, Israel, for running seed and leaf extracts by LC-MS and Änne Michaelis and Saleh Alseekh, Max Planck Institute of Molecular Plant Physiology (MPI-MP), Germany, for running LC-MS of SALK mutant lines. We would like to thank Si Wu, University of Stanford, USA, and Alvaro Cuadros-Inostroza, MetaSysX, Germany, for providing introduction and scripts for GWAS analysis. We thank Marcin Luzarowski for his help in running Genedata. Furthermore, we would like to acknowledge the valuable input of Joachim Kopka and Mark Stitt, MPI-MP, Germany, for this project. We thank the members of the green team at the MPI-MP for their help in growing the Arabidopsis accessions. We thank Josef Bergstein for taking pictures of the Arabidopsis plants. T.N. acknowledges the support by the IMPRS-PMPG program.

## Supplementary Figures

**Supplementary Figure S1:**
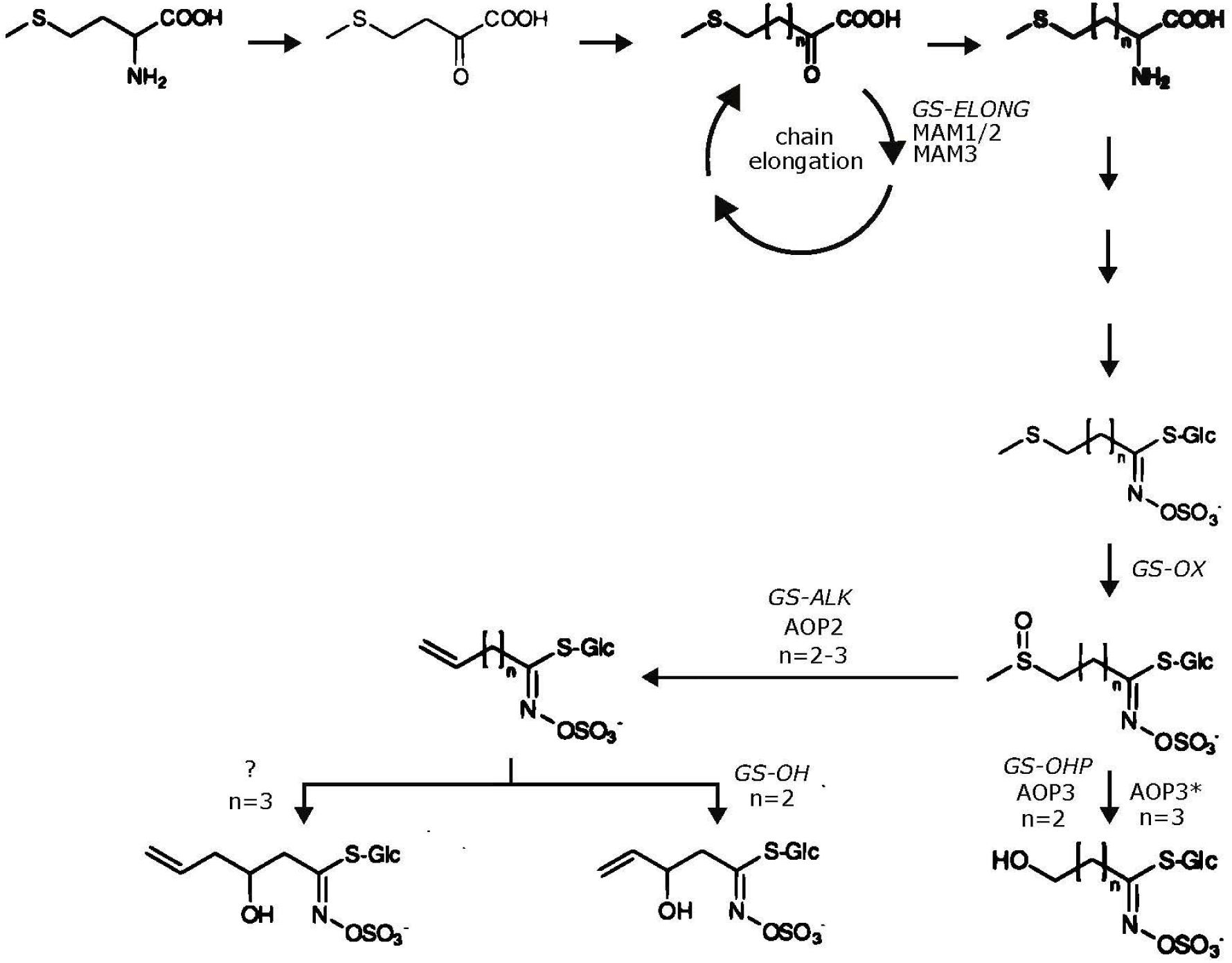
Simplified pathway for aliphatic glucosinolates. The biosynthetic pathway starts with the deamination of methionine to an 2-oxo acid by a branched-chain amino acid aminotransferase. Subsequently, the 2-oxo acid enters a cycle of three successive transformations: condensation with acetyl-CoA by MAM, isomerization by an isopropylmalate isomerase (not shown), and oxidative decarboxylation by an isopropylmalate dehydrogenase (not shown, Sønderby et al., 2010). One round of chain elongation leads to an elongation of one ethylene group (-CH_2_-). After several intermediate steps, including sulfur incorporation, the side chain of aliphatic glucosinolates is modified by *GS-OX*, *GS-ALK*, *GS-OH*, and *GS-OHP* (Sønderby et al., 2010). The biosynthetic pathway is modified from Sønderby et al. (2010). AOP: alkenyl hydroxyalkyl producing; MAM: methylthioalkylmalate synthase; *: predicted enzyme; ?: unknown enzyme.

**Supplementary Figure S2:**
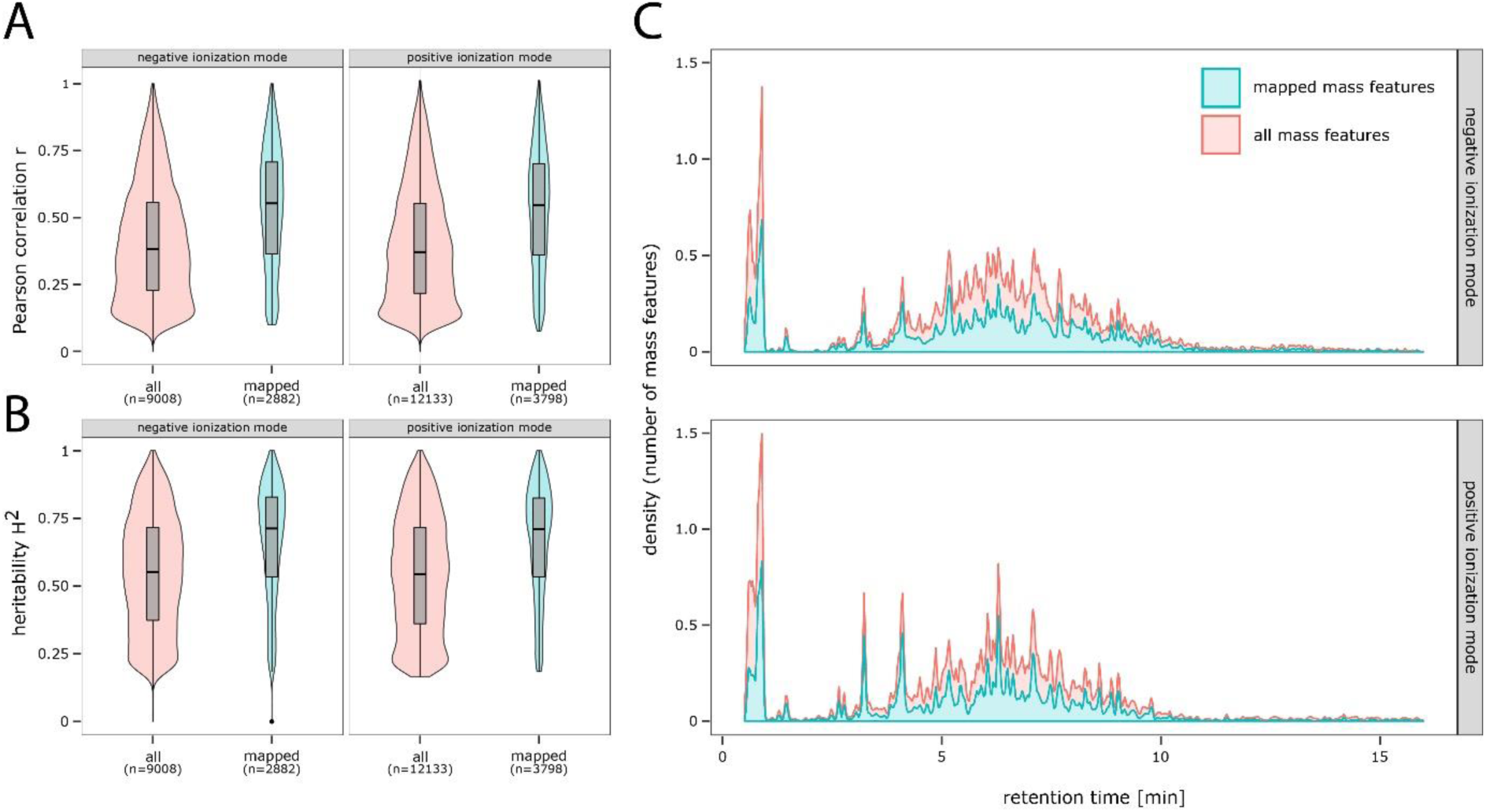
Metabolite data sets for replicate 1 and 2 (genome-wide association studies). A: Pearson correlation coefficient (*r*) values of intensity values for matched mass feature pairs for negative and positive ionisation mode. *r* values of mapped mass features were higher compared to the random pairs of the core set. B: Broad-sense heritability (H^2^) of intensity values for matched mass feature pairs for negative and positive ionisation mode. H^2^ values of mapped mass features were higher compared to the random pairs of the core set. C: Distribution along retention time for all matched mass feature pairs and those that were mapped to at least one locus with LOD ≥ 5.3 for negative and positive ionisation mode. LOD: logarithm of odds.

**Supplementary Figure S3:**
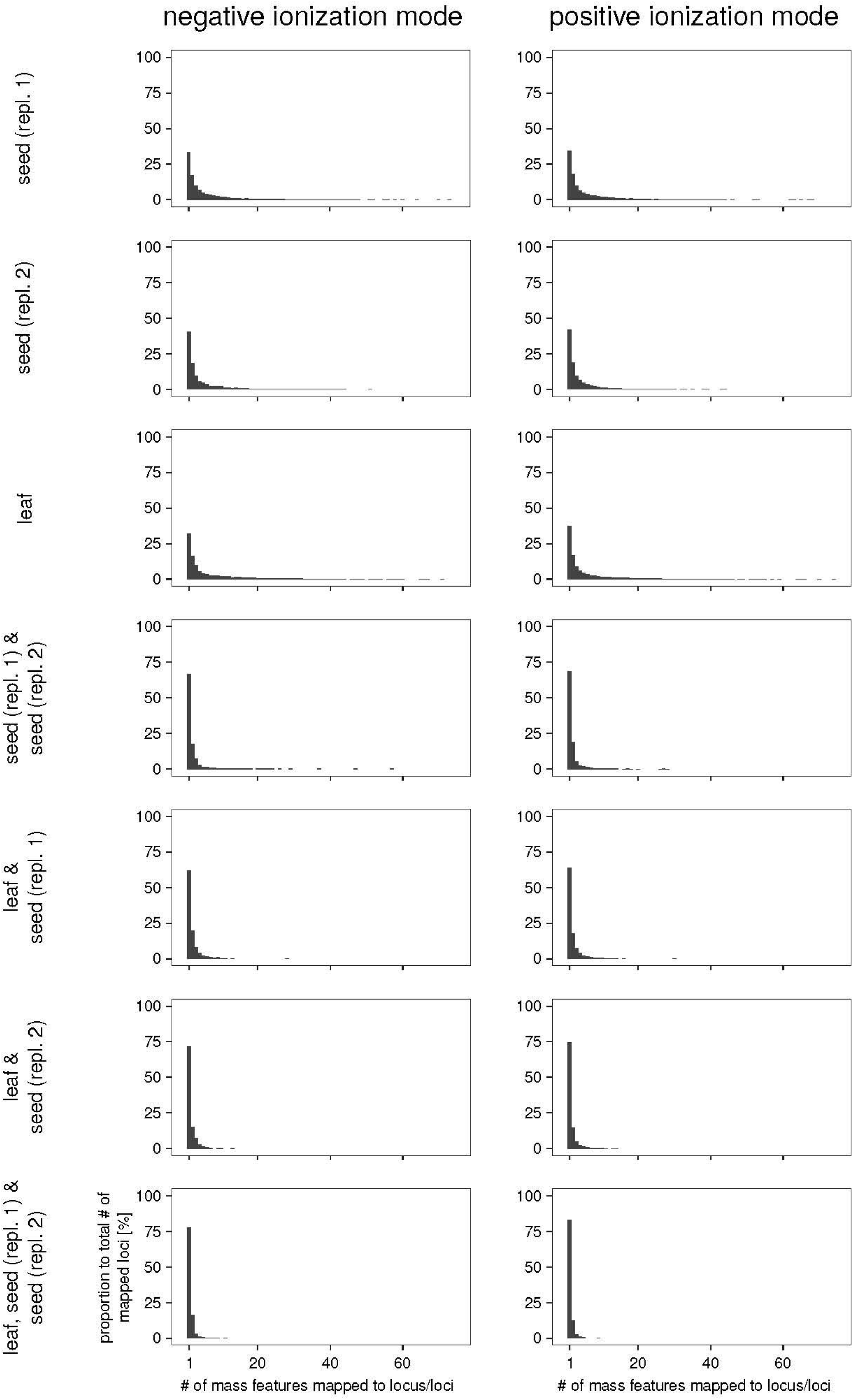
Distribution of number of mass features mapped to locus/loci per intersection set. Each panel shows how often a specific mass feature was mapped to a locus/loci per intersection set (e.g., a value of 1 on the x-axis means that this mass feature was mapped to only one locus). For the intersection sets seed (replicate 1), seed (replicate 2), and leaf a higher proportion of mass features was found that were associated with several loci. This can be attributed to random sources of measurement errors, to associations of non-causative markers with a given trait, to linkage with causative markers (Korte and Farlow, 2013), or to environmental effects. The mass features of the other intersection sets show an association with mainly one locus.

**Supplementary Figure S4:**
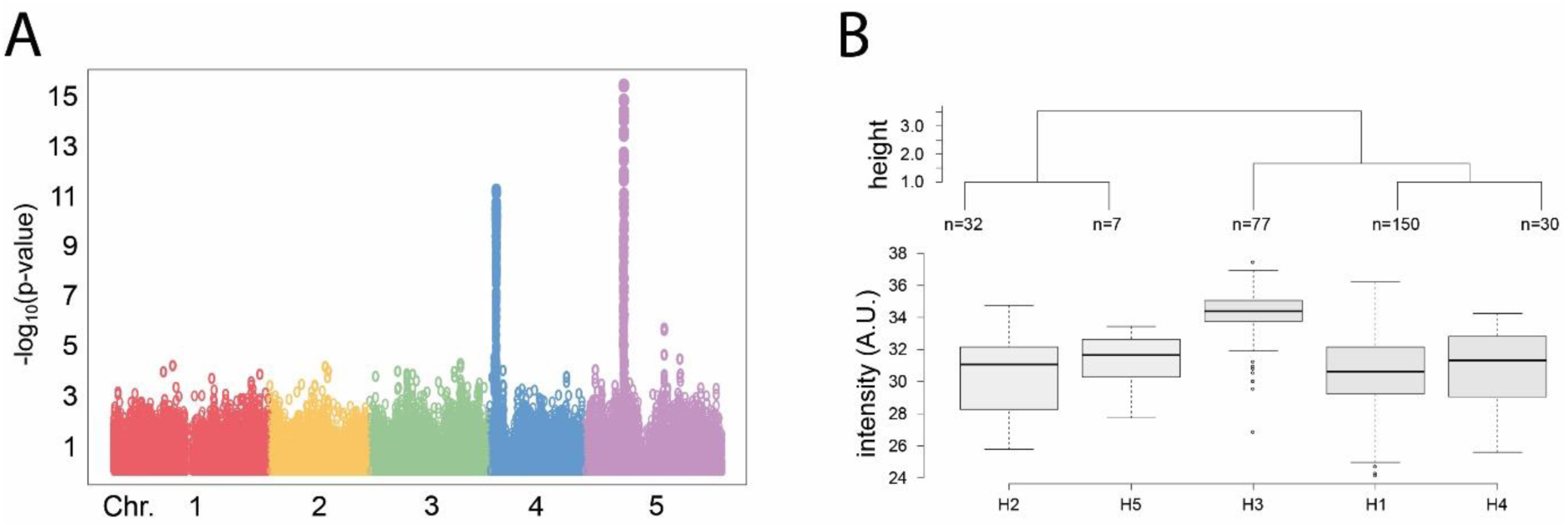
Genome-wide association mapping for 3-hydroxypropyl glucosinolate (negative ionization mode). A: The Manhattan plot of 3-hydroxypropyl glucosinolate shows two peaks in each replicate on chromosomes 4 (highest LOD: 11.22) and 5 (15.39). These loci contain the genes *AOP1, AOP2*, and *AOP-3* (chromosome 4), *MAM1* and *MAM3* (chromosome 5) that are involved in glucosinolate biosynthesis. B: Haplotype analysis of metabolite levels of 3-hydroxypropyl glucosinolate. The nucleotide sequence differences were statistically associated with the levels of 3-hydroxypropyl glucosinolate (ANOVA q-value: 7.05e-21 for replicate 2). Only data for replicate 2 is shown in A and B. The data for replicate 1 is depicted in Figure 2. *AOP*: *alkenyl hydroxyalkyl producing*; A.U.: arbitrary units; LD: linkage disequilibrium; LOD: logarithm of odds; *MAM*: *methylthioalkylmalate synthase*.

**Supplementary Figure S5:**
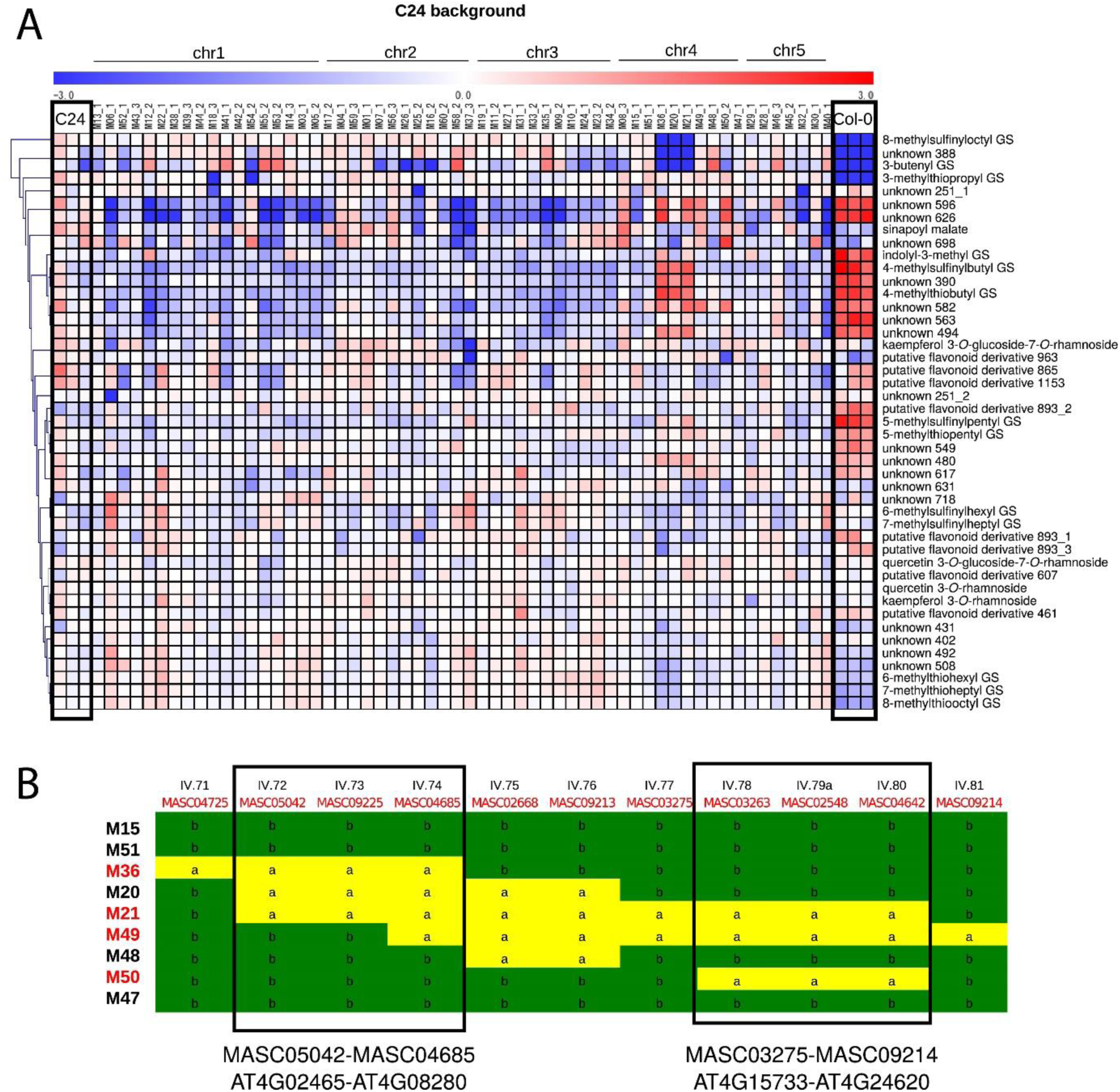
QTL mapping using near-isogenic introgression lines between C24 and Col-0 (C24 background). A: Heatmap of relative seed metabolite levels for C24, Col-0 and NILs with C24 background. B: Genomic region for NILs and allele identity based on MASC genomic markers. The highlighted regions refer to the lines with Col-0 alleles that show altered levels of glucosinolates and of the unknowns 596 and 626. a: genomic region from Col-0; b: genomic region from C24; GS: glucosinolates; NIL: near-isogenic line.

**Supplementary Figure S6:**
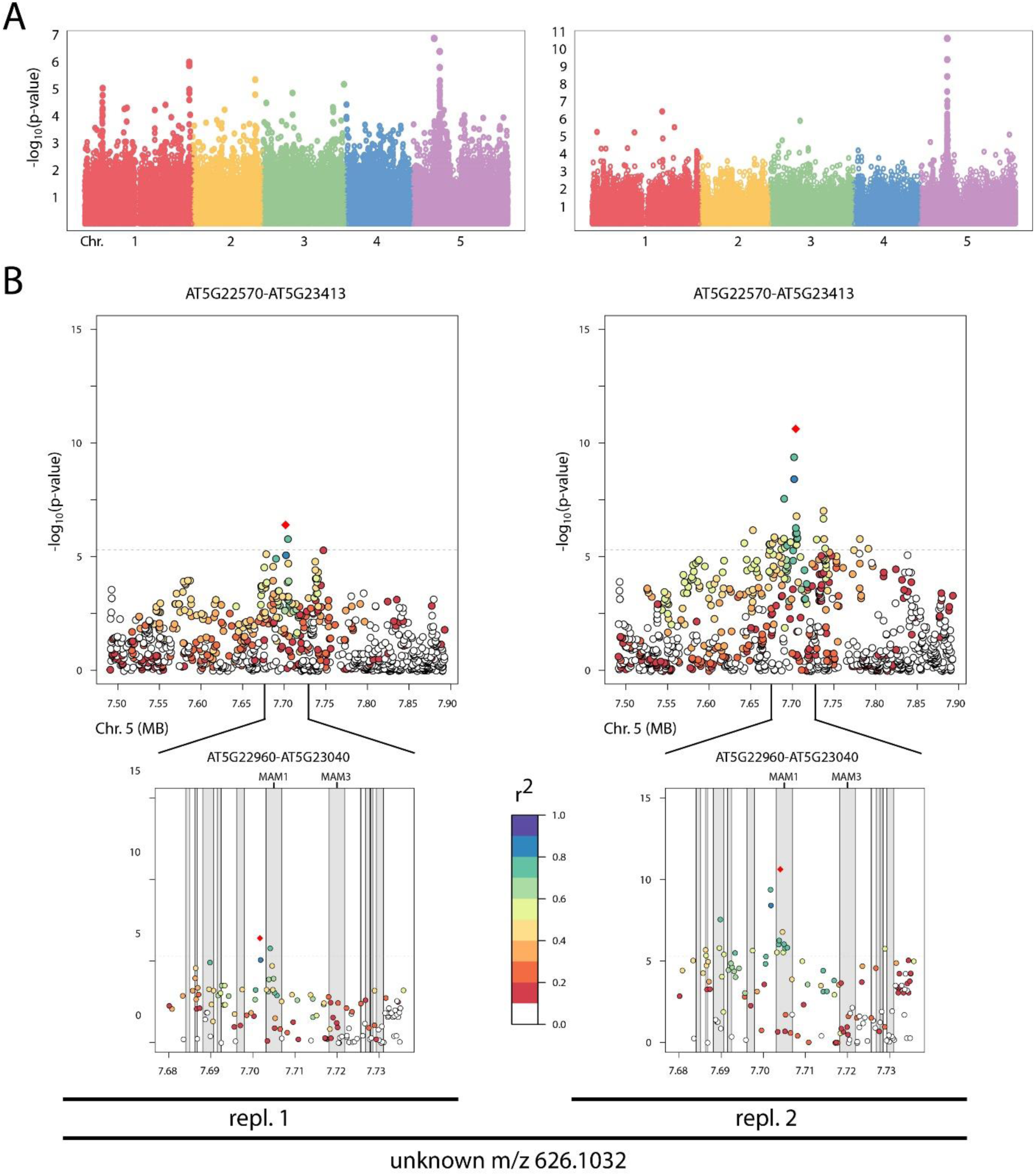
Manhattan plot and linkage disequilibrium analysis of the unknown 626 (negative ionization mode). A: Manhattan plot for replicate 1 (left) and replicate 2 (right). The plots show an association of the unknown 626 with a genomic region on chromosome 5 containing *MAM1* and *MAM3*. B: The highest LOD score is obtained within the region of *MAM1*. The LOD values decay quickly when moving away from *MAM1*. LOD: logarithm of odds; MAM: methylthioalkylmalate synthase.

**Supplementary Figure S7:**
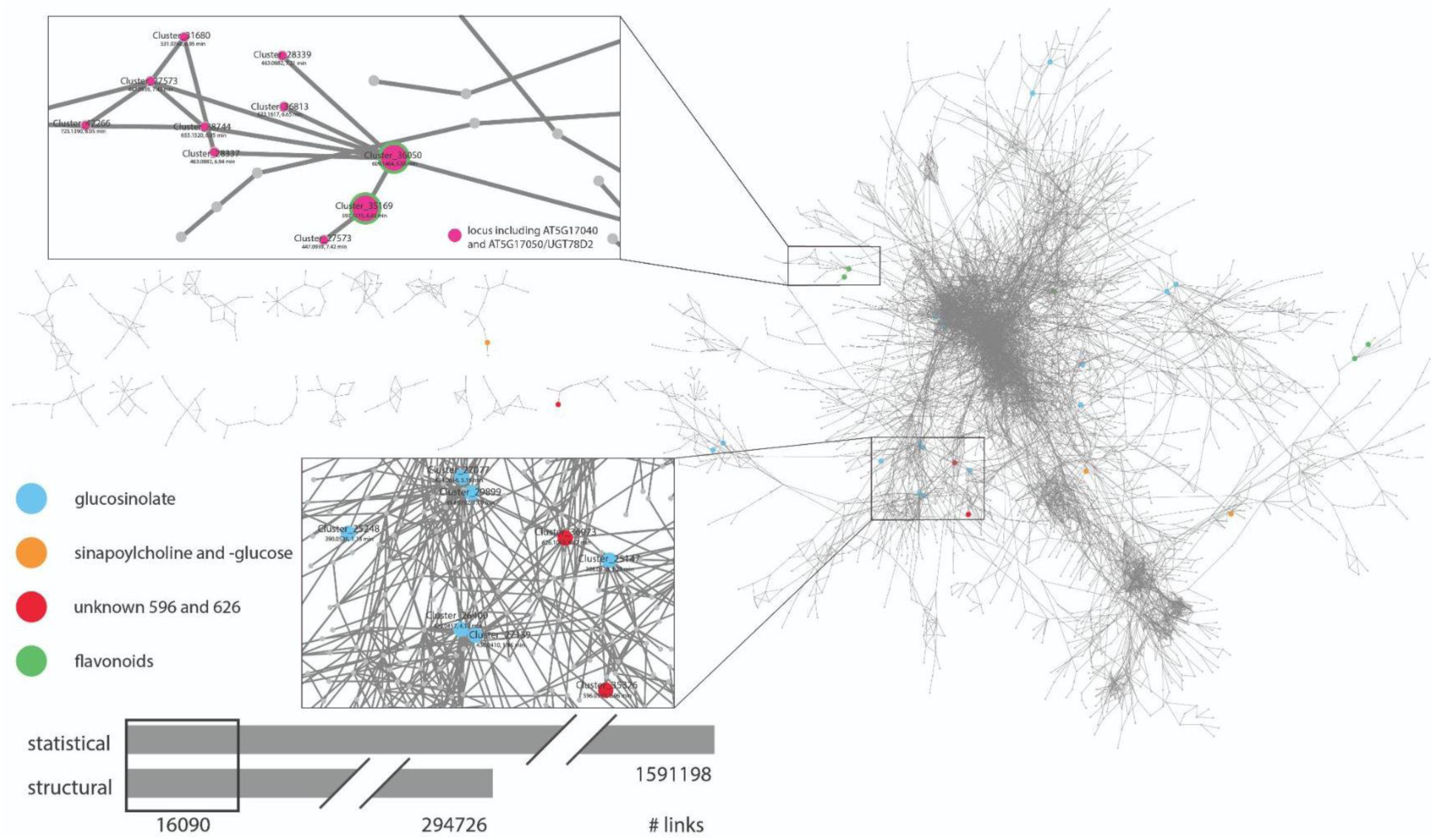
Metabolite network of mapped mass features (negative mode). The network was constructed by `MetNet` and consists of 8819 mass features (vertices) and 16090 joint edges from the structural and statistical network inference. The 8819 mass features showed associations to at least one locus in one replicate. Mass features corresponding to annotated metabolites are highlighted (glucosinolates, sinapoylcholine and -glucose, unknown 626, and flavonoids). The figure shows only the major network components. *m/z* and retention time values are taken from the alignment of mass spectrometric data of replicate 1. Linking mass features to the unknown 626 showed association with the *GS-ALK*/*GS-OHP*, *and*/*or GS-ELONG* loci with a LOD > 5.3. Linking mass features to quercetin-containing flavonols showed association with the locus containing *AT5G17040* and *AT5G17050*/*UGT78D2* with a LOD > 5.3. The highlighted mass features also showed associations with other loci (LOD > 5.3). LOD: logarithm of odds.

**Supplementary Figure S8:**
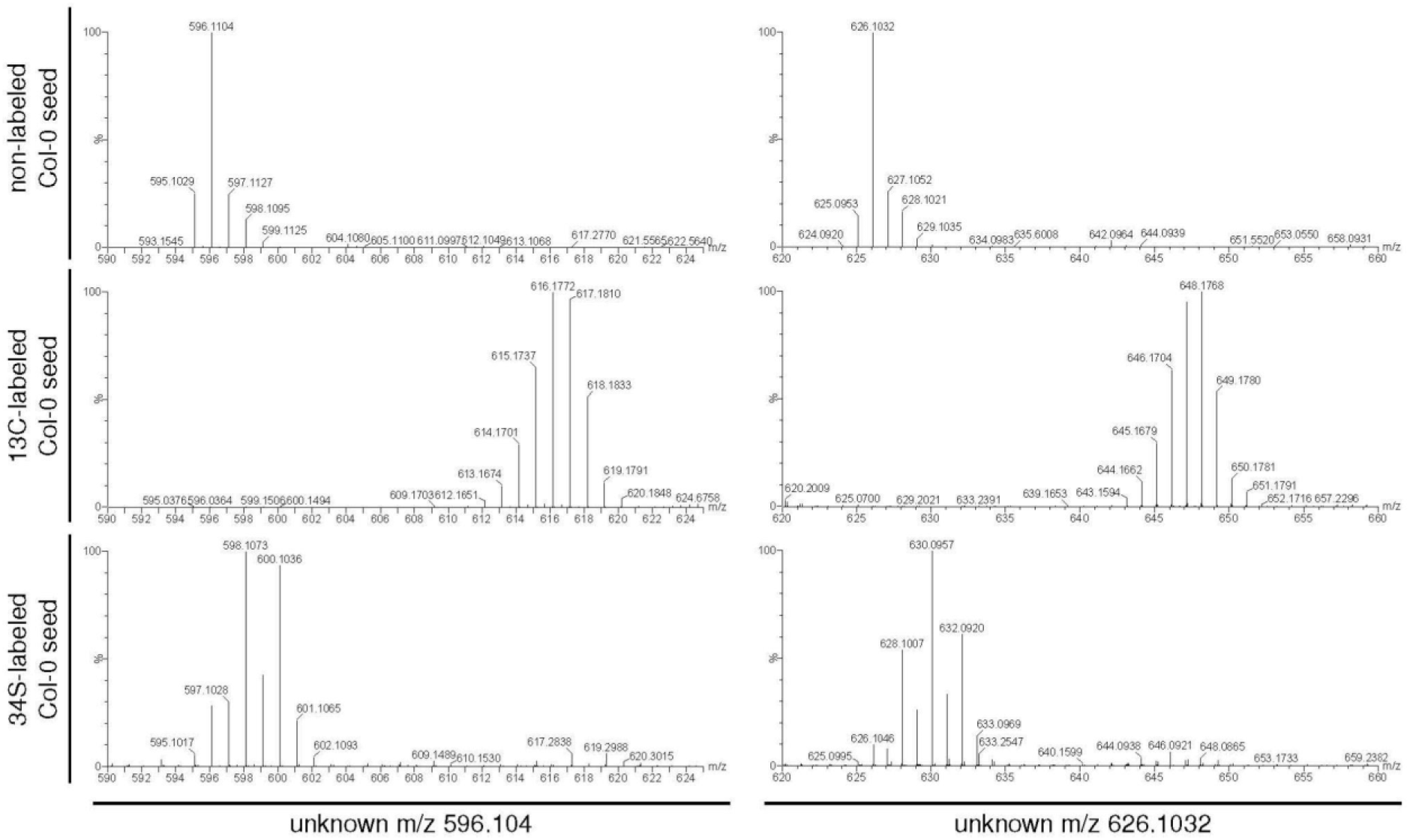
Effects of ^13^C and ^34^S feeding on the *m/z* of the unknowns 596 and 626 (negative ionization mode). The measurement was performed via LC-QTOF-MS and revealed that the unknown 596 (*m/z* 596.1104) and 626 (m/z 626.1032) contains 20 C atoms and 22 C atoms based on isotope feeding with ^13^C, respectively, and two S atoms based on isotope feeding with ^34^S.

**Supplementary Figure S9:**
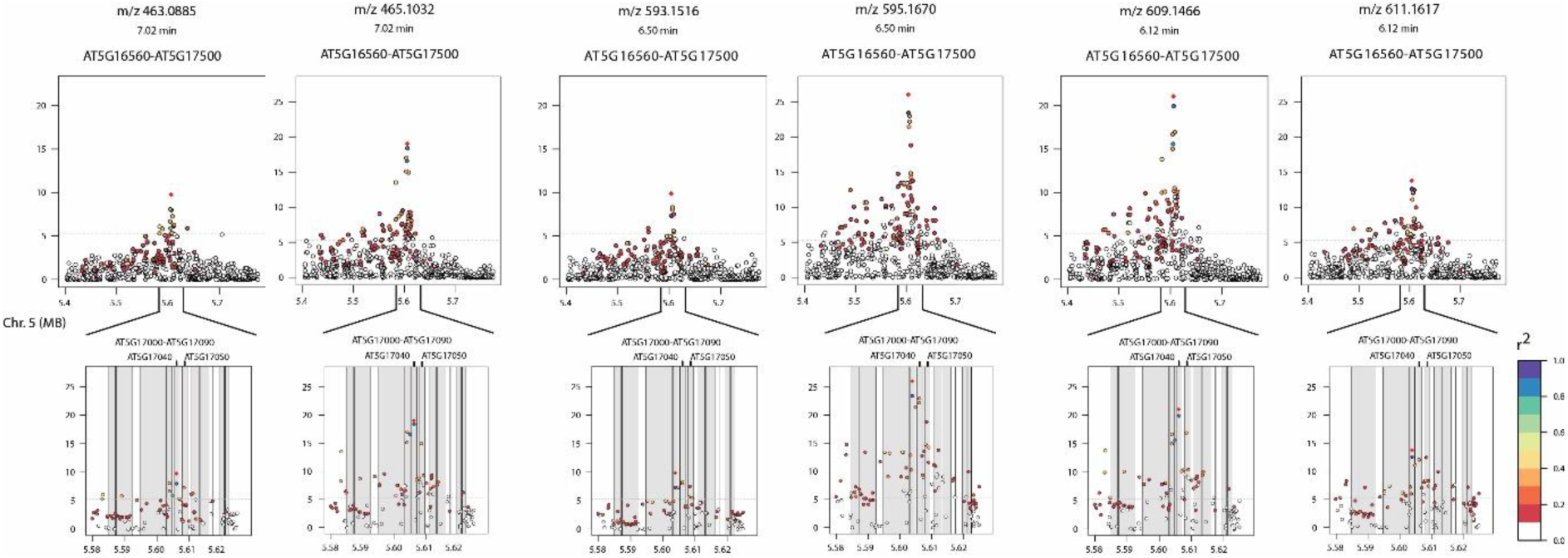
Flavonoid biosynthetic pathway: Linkage disequilibrium analysis for quercetin-containing flavonols (negative and positive ionization mode). The highest LOD is achieved for a SNP within the region of *AT5G17030* or *AT5G17040*. Standardized LD r^2^ is relatively low for the SNPs that are located within the gene *AT5G17050*. Only data for replicate 2 is shown. The data for replicate 1 is depicted in Figure 3. LD: linkage disequilibrium; LOD: logarithm of odds; MB: megabase; SNP: single nucleotide polymorphism.

**Supplementary Figure S10:**
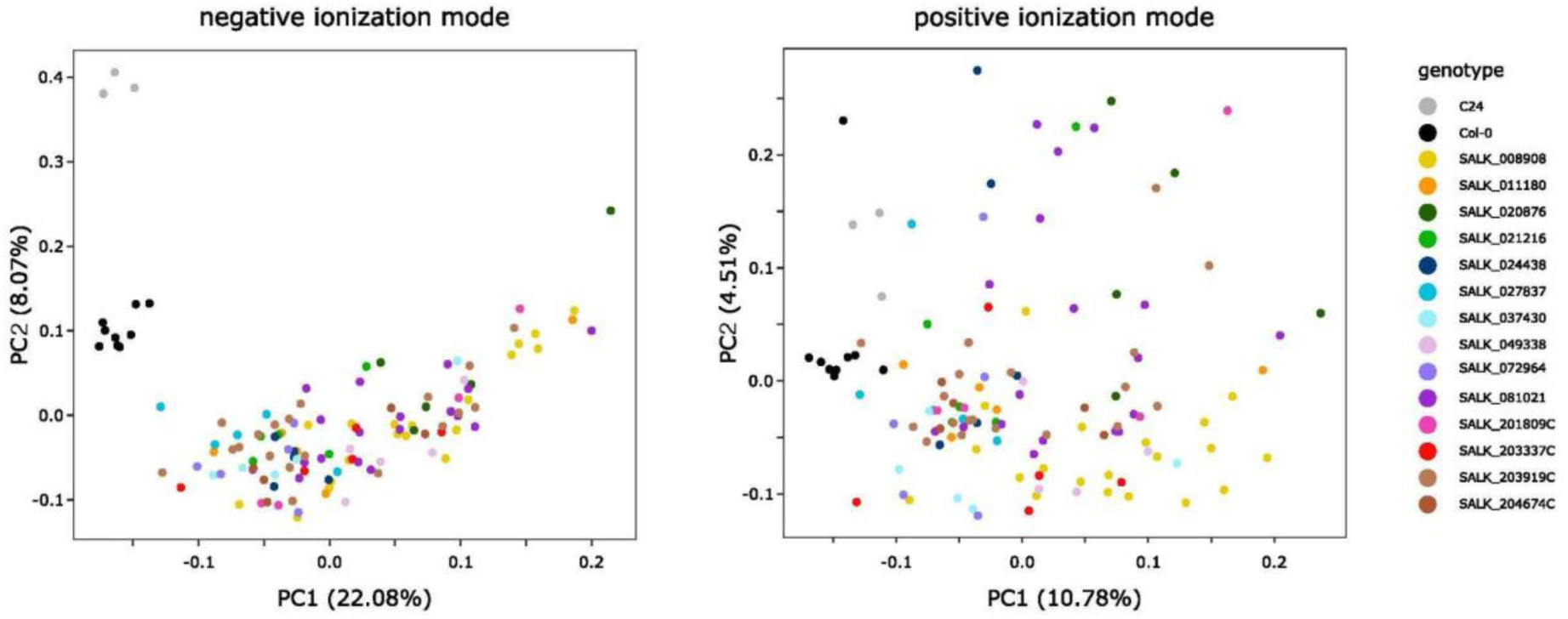
Principal component analysis for metabolite analysis of SALK lines (replicate 1). Negative ionization mode: The first two PCs explain 30.15% of the variance in the seed data set. 27716 mass features were used for the PC analysis. The projection of the mass feature levels of C24 (gray dots) and Col-0 (black dots) are located close to each other for each genotype and are distinct from the SALK mutants. Positive ionization mode: The first two PCs explain 15.29% of the variance in the seed data set. 28750 mass features were used for the PC analysis. The projections of the mass feature levels of C24 (gray dots) and Col-0 (black dots) are located close to each other for each genotype. PC1 and PC2 do not separate C24 and Col-0 from the SALK line mutants. The PCA for the replicate 2 is not shown here. PC: principal component.

**Supplementary Figure S11:**
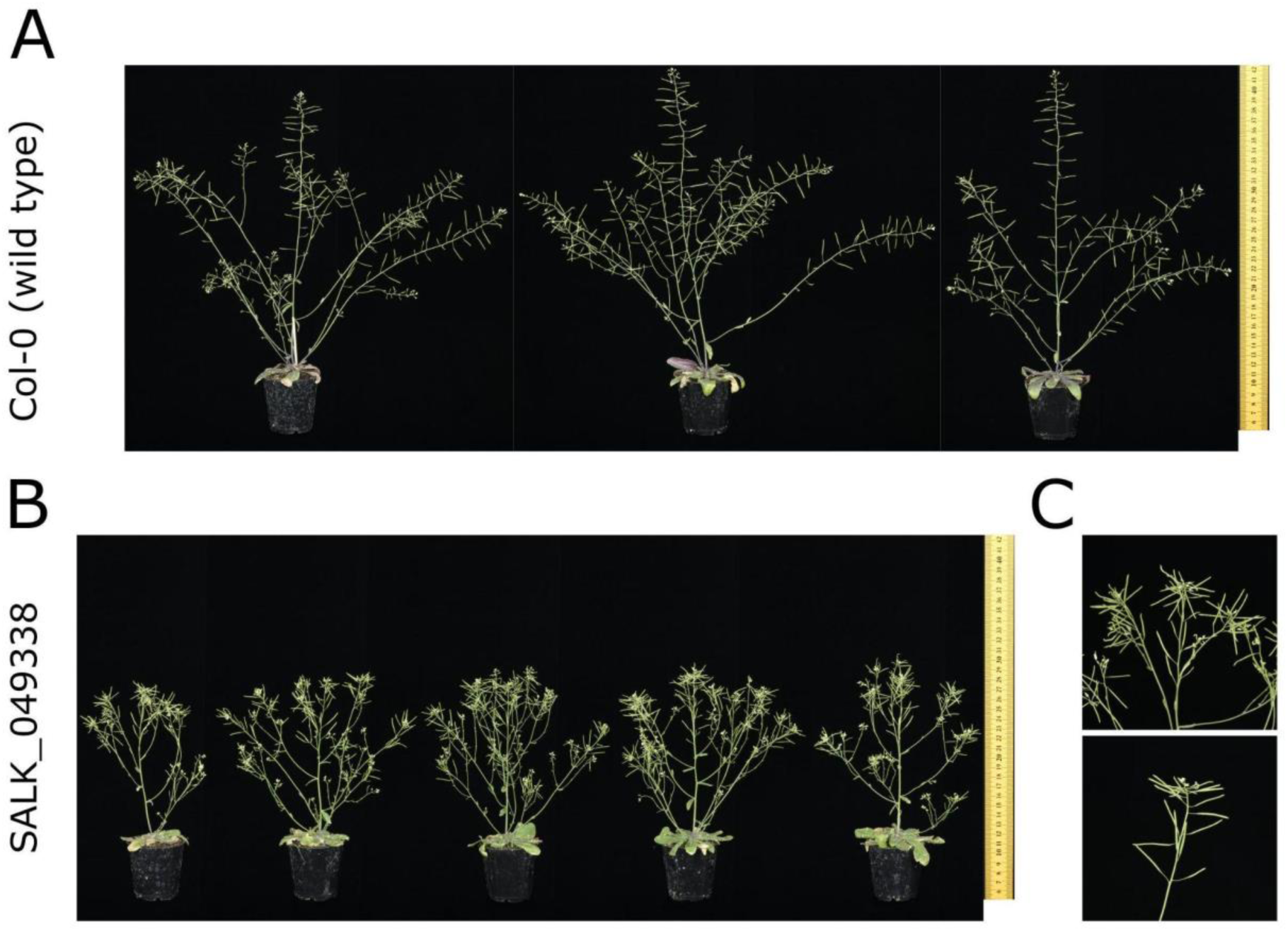
Phenotype of Col-0 and SALK_049338 mutant lines. A: Three biological replicates of wild type Col-0. B: Five biological replicates of SALK_049338 mutant lines, referring to a T-DNA insertion in the exon of *AT5G17050* (*UGT78D2*). The lines exhibited a dwarf phenotype with a loss of apical dominance as reported previously by Yin et al. (2014). C: Detail of inflorescence of line SALK_049338.

